# Modeling the zebrafish gut microbiome’s resistance and sensitivity to climate change and parasite infection

**DOI:** 10.1101/2025.03.28.644597

**Authors:** Michael J. Sieler, Colleen E. Al-Samarrie, Kristin D. Kasschau, Mike L. Kent, Thomas J. Sharpton

## Abstract

As climate change increases global water temperatures, ecologists expect intestinal helminth infection ranges to expand and increase the health burden on aquatic organisms. However, the gut microbiome can interact with these parasites to influence infection outcomes, raising the possibility that its response to increasing temperatures may help buffer against increased infection burden. To evaluate this hypothesis, we sought to determine if the microbiome is resistant or resilient to the stressors of increased water temperature, helminth exposure, and their combination, and whether this variation linked to infection outcomes. We leveraged the zebrafish (*Danio rerio*) model organism to measure how these variables relate to the temporal dynamics of the gut microbiome. In particular, we exposed adult zebrafish to the parasitic whipworm Pseudocapillaria tomentosa across three different water temperatures (28°C, 32°C, 35°C), and analyzed fecal microbiome samples at five time points across 42 days. Our findings show that parasite exposure and water temperature independently alter gut microbiome diversity. Moreover, we find that water temperature moderates the association between parasite infection and the gut microbiome. Consistent with this observation, but at odds with current expectations, we find that increasing water temperature reduces parasitic infection in fish. Overall, our results indicate that water temperature alters the contextual landscape of the gut microbiome to impact its response to an exogenous stressor of an intestinal parasite in zebrafish. Furthermore, our findings represent the first report of the effects of elevated temperature on parasitic nematode development in a fish host. Importantly, our study demonstrates that climate change may have unanticipated and environmentally contingent impacts to vertebrate gut microbiomes and health outcomes in response to an exogenous stressor.

## Introduction

The steady increase in global temperatures due to climate change challenges vertebrate health (1). These threats to vertebrate health take on many forms, including the expected expansion of infectious agents (2,3). Of particular concern are the increased infection burdens faced by aquatic organisms experiencing increasing water temperatures (4). Due in part to the varied coincident effects of climate, the impacts of a warming climate on aquatic organisms are anticipated to be nonuniform in effect (4,5) and vary biogeographically (4,6), which in turn complicates harm mitigation and conservation efforts (7). Consequently, there’s an urgent need to better understand climate change’s contextual impacts on organisms depending on the unique environmental conditions of the ecosystems they inhabit.

In recent years researchers have considered that climate change may also elicit harm to vertebrates by disrupting the composition of their gut microbiome (8). While prior work has shown that varying temperatures impacts gut microbiome composition across a variety of vertebrate host species (9), less is known about how coincident variables, such as parasite or pathogen exposure, collide with temperature to drive variation in the gut microbiome. These potential interaction effects are important to quantify, because it may be that they elicit even greater effects on the gut microbiome than anticipated by investigations of temperature alone, and could possibly result in increased frequency of dysbiotic disorders. It’s important to elucidate these interactions because increasing work points to the gut microbiome as a key determinant of whether vertebrate physiology is able to buffer against stress (9,10), and whether temperature induced perturbations to the gut microbiome may sensitize individuals to subsequent stressors (11).

To answer these questions, we evaluated the gut microbiome’s temporal response to an exogenous stressor across a gradient of environmental conditions. To do so, we levered the zebrafish (*Danio rerio*) model organism to measure how gut microbiomes differ across fish reared to adulthood at one of three water temperature conditions (28°C, 32°C, or 35°C; Fig. 1). Zebrafish are highly thermal tolerant, capable of existing across a wide spectrum of temperature ranges from 4°C to 40°C (12), but living outside their thermal optimum can come at a physiological and microbial cost (12,13). While much is known about the thermal range of zebrafish, the effects of altered water temperature on their gut microbiome structure has not been elucidated. We also sought to determine if water temperature affected how the microbiome and host responds to exposure to and infection by intestinal nematode *Pseudocapillaria tomentosa*, a common source of disease in zebrafish facilities (14). *P. tomentosa* is known to cause high mortality and disrupts the gut microbiome (14,15). Yet, it is not known if or how water temperatures mediates this interaction. Overall, our study sought to clarify the environmentally dependent context of a gut microbiome’s resistance and sensitivity to climate change-relevant stressors.

**Figure 1.**
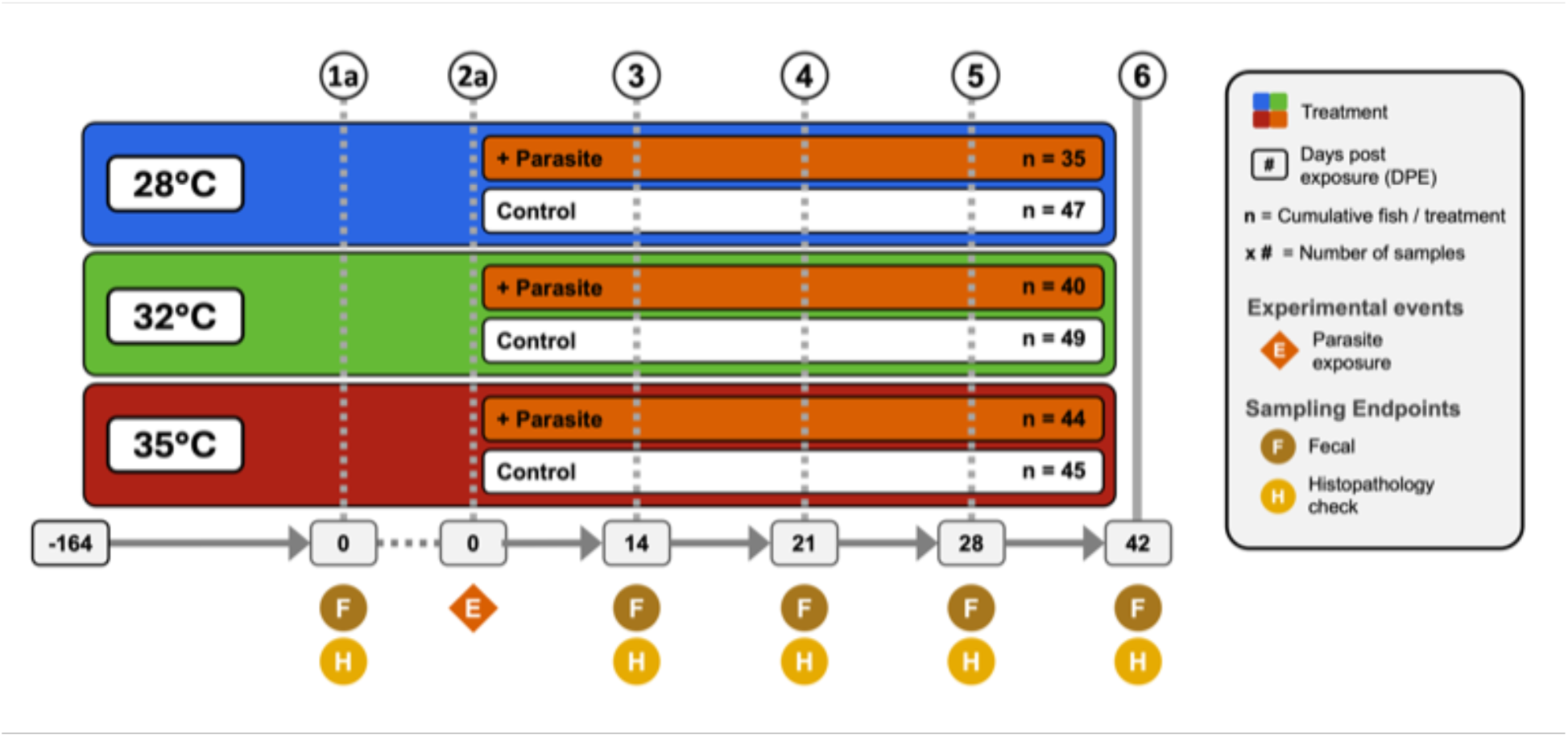
Experimental design showing treatments and husbandry events during the course of the study. Symbols indicate when an experimental event occurred at each time point. (1) 260 fish were assigned and acclimated to one of three water temperature groups (e.g., 28°C, 32°C, or 35°C) and reared from 0 to 164 days post fertilization (dpf). (2a) At 164 dpf (or 0 days post exposure; DPE), fecal collections were collected from a random selection of five fish per tank (n = 60). Additionally, histological and wet mount assessments were conducted on selected fish to assess presence of infection and infection burden. (1b) Afterwards, a cohort of fish from each water temperature group were exposed to *Pseudocapillaria tomentosa*. (3-6) Subsequent fecal samples were collected and histopathological assessments were conducted at 14 dpe (n = 54), (3) 21 dpe (n = 48), (4) 28 dpe (n = 47), and (5) 42 dpe (n = 51).

## Results

### Water temperature shapes gut microbiome structure

To determine how zebrafish reared across a gradient of increasing water temperatures impacts the structure of the gut microbiome, we reared 260 zebrafish across one of three water temperatures (28°C, 32°C or 35°C) until 206 days-post fertilization (dpf; Fig. 1). Additionally, within each temperature cohort, fish were evenly divided into two additional treatment groups: either unexposed or exposed to the intestinal helminthic parasite *Pseudocapillaria tomentosa*. Microbiome samples were collected at five time points between 164 and 206 dpf. In the parasite exposed cohort, fish were exposed to *P. tomentosa* following microbiome sampling at 164 dpf, or 0 days post exposure (dpe). Four subsequent microbiome samples were collected at 14 dpe (178 dpf), 21 dpe (185 dpf), 28 dpe (192 dpf), and 42 dpe (206 dpf). Within the parasite unexposed fish cohort, we built generalized linear models (GLM) to determine if water temperature associated with variation in one of four measures of alpha-diversity: Simpson’s Index, Shannon Entropy, richness, and phylogenetic diversity (Table S2A.1). An ANOVA test of these GLMs revealed that alpha-diversity varied as a function of temperature for all measures (P<0.05; Fig. 2A; Table S2A.2), except Shannon Entropy (P>0.05; Table S2A.2). A post hoc Tukey test clarified that alpha-diversity scores did not significantly differ between 28°C and 32°C water temperature reared fish for each diversity metric we measured (P>0.05; Table S2A.3). However, we observed significant differences in diversity between 28°C and 35°C water temperature reared fish across Simpson’s Index, richness and phylogenetic alpha-diversity measures (P<0.05; Table S2A.2), and between 32°C and 35°C water temperature reared fish as measured by richness and phylogenetic diversity metrics (P<0.05; Table S2A.2). These results indicate that water temperature associates with fish gut microbiome diversity, and that water temperature may differentially impact particular microbial clades of the gut.

**Figure 2.**
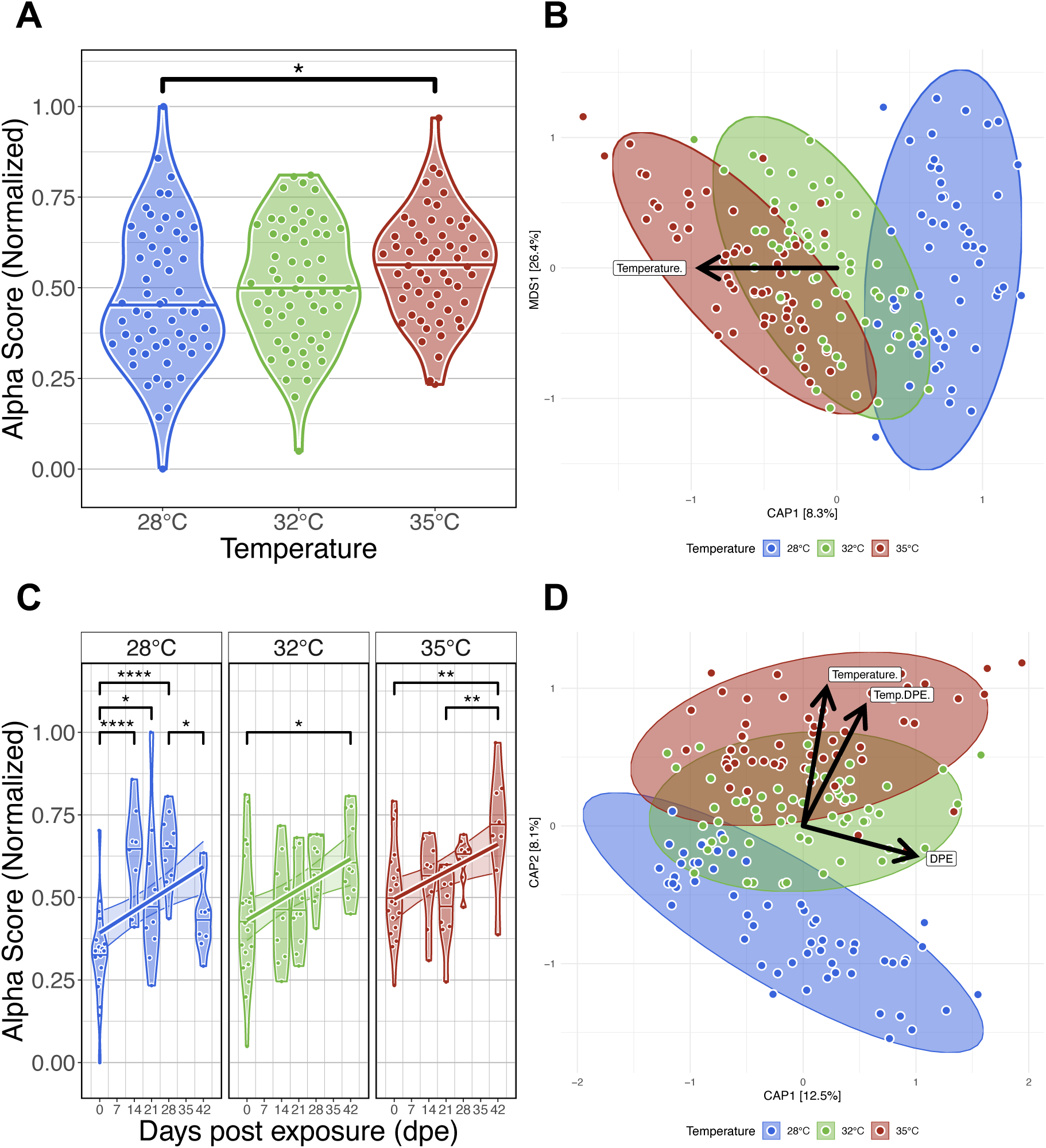
Effects of water temperature on zebrafish gut microbiomes. (A) Simpson’s Index of diversity shows that gut microbiome diversity significantly differs between fish reared at 28°C and 35°C water temperatures. (B) Capscale ordination based on the Bray-Curtis dissimilarity of gut microbiome composition constrained on the main effect of temperature. The analysis shows that gut microbiome composition significantly differs between fish reared at different water temperatures. (C) Simpson’s Index of diversity shows microbial gut diversity increases with time from 0 days post exposure (dpe) to 42 dpe, irrespective of water temperature. (D) Capscale ordination of gut microbiome composition based on the Bray-Curtis dissimilarity constrained on the main effects of water temperature and time (days post exposure, dpe), and their interaction. The analysis shows that shows that gut microbiome composition differs between fish across time depending on water temperature. Ribbons and ellipses indicate 95% confidence interval. Only statistically significant relationships are shown. A “*” indicates statistical significance below the “0.05” level. Black arrows indicate direction of greatest change in the indicated by covariates.

To evaluate how temperature associates with microbiome composition in parasite unexposed fish, we quantified dissimilarity amongst all samples and generated distance matrices using the Bray-Curtis, Canberra and half-weighted UniFrac distance metrics. Using permutational multivariate analysis of variance (PERMANOVA), we assessed whether increasing water temperatures explained variance in gut microbial community composition. A PERMANOVA test indicated that microbial communities were significantly stratified by water temperature as measured by all beta-diversity metrics (PERMANOVA, P<0.05; Fig. 2B; Table S2B.1). These results indicate that microbial communities of fish reared at the same water temperature are more consistent in composition to one another than fish reared at different water temperatures. Additionally, we assessed beta-dispersion, a measure of variance, in the gut microbiome community compositions for each water temperature group. We find the beta-dispersion levels did not significantly differ between the water temperature groups (P>0.05; Table S2B.2). These results indicate that fish reared at different water temperatures are consistent in community composition.

Next, we compared our results across five time points between 0- and 42 dpe to determine how water temperature impacts the successional trajectories of gut microbiome diversity and composition. Linear regression revealed gut microbial alpha-diversity was significantly associated with the main effect of time for each alpha-diversity metric we assessed (P<0.05; Fig. 2C; Table S2C.1-2). Moreover, we found a temperature dependent effect on time as measured by richness and phylogenetic diversity metrics (P<0.05; Table S2C.1-2). A post hoc Tukey test clarified that microbiome diversity significantly differed between 0- and 42 dpe fish reared at 28°C as measured by richness and phylogenetic diversity (P<0.05; Table S2C.3), between 0- and 42 dpe fish reared at 32°C as measured by all alpha-diversity metrics (P<0.05; Table S2C.3), and between 0- and 42 dpe fish reared at 35°C as measured by Shannon Entropy and Simpson’s Index (P<0.05; Table S2C.3). These results indicate that gut microbial alpha-diversity increases over time irrespective of water temperature.

A PERMANOVA test detected significant clustering of microbial gut community composition based on the interaction of water temperature and time as measured by all beta-diversity metrics (PERMANOVA, P<0.05; Fig. 2D; Table S2D.1). These results indicate that microbial communities of fish reared at the same water temperature are more consistent in composition to one another across time than fish reared at different water temperatures. Moreover, a pairwise analysis of beta-dispersion found significantly elevated levels of dispersion between fish reared across different temperatures and time as measured by all beta-diversity metrics (P<0.05; Table S2D.2). These results indicate that gut microbial community composition varies inconsistently between water temperature groups in a time-dependent manner. Collectively, these results indicate that zebrafish gut microbiomes communities stratify by temperature, and the trajectory of gut microbiome successional development varies depending on water temperature.

### Infection burden is highest in fish reared at lower water temperatures

Next, we evaluated infection outcomes of zebrafish reared at different water temperatures and exposed to the intestinal helminthic parasite *Pseudocapillaria tomentosa*. To determine whether water temperature affects infection burden, we exposed zebrafish to 50 *P. tomentosa* eggs per liter of tank water at 164 days post-fertilization (dpf). Infection outcomes were assessed using wet mount and histological evaluation at 0, 14, 21, 28, and 42 days post-exposure (dpe). We built a negative binomial general linear model to compare infection burden (total worm counts) between fish reared at different water temperatures (Table S3B.1). The regression analysis found a statistically significant effect of temperature on infection burden (P < 0.05; Fig. 3B; Table S3B. 2). However, we did not find a statistically significant interaction effect between water temperature and time on infection burden (*P* > 0.05; Table S3B.3).

**Figure 3.**
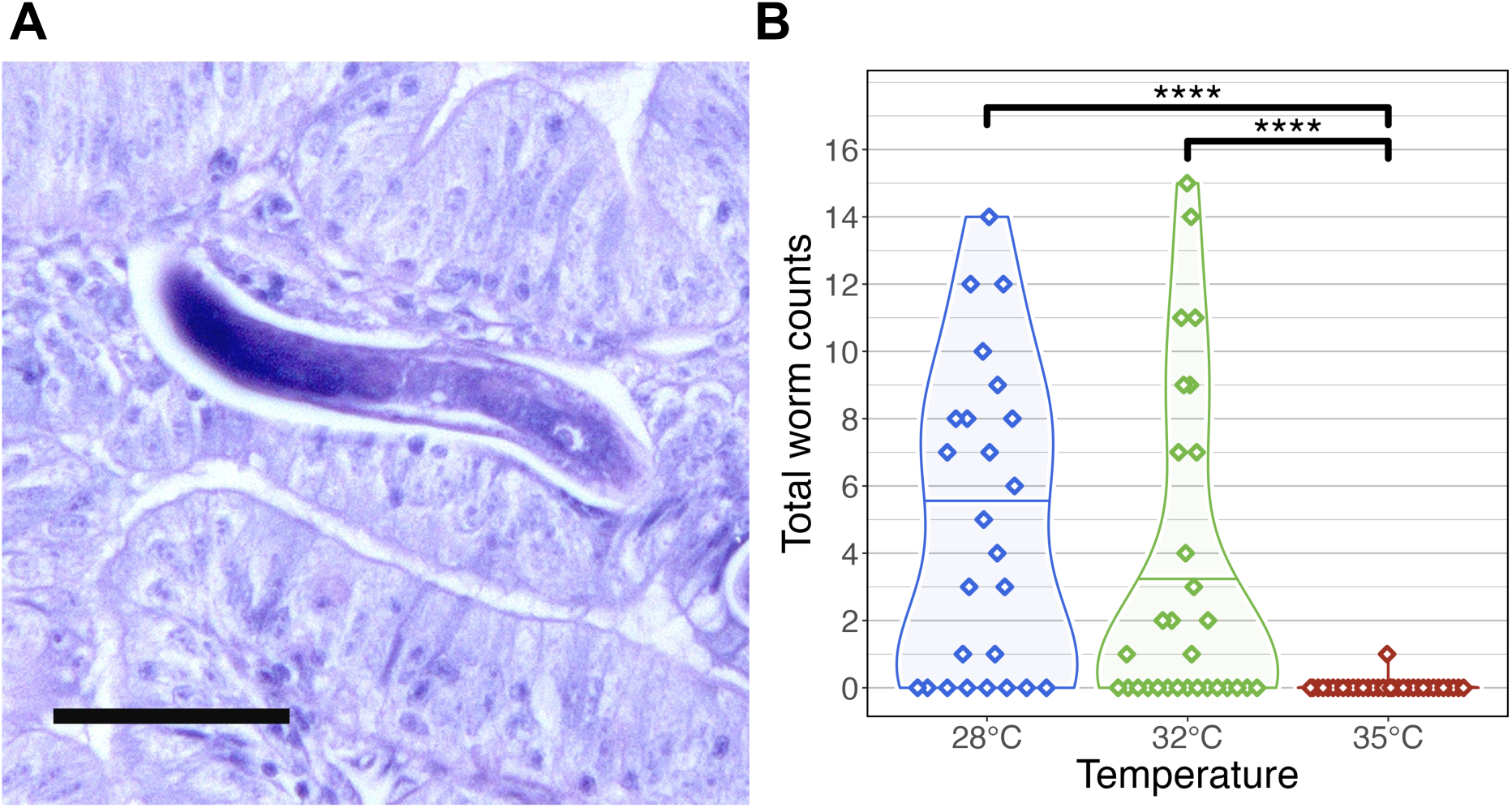
Infection outcomes in zebrafish exposed to *Pseudocapillaria tomentosa*. (A) Histological sections stained with H&E stain in zebrafish exposed to *P. tomentosa* examined at 35°C at 21 days post exposure. Arrow = larval worms, sagittal and cross sections. Bar = 50 µm (B) Infection outcome analysis of fish exposed to *P. tomentosa* (n = 89) by temperature. Fish reared at 28°C and 32°C water temperatures had significantly different infection burden to fish reared at 35°C water temperature. Only one fish in our microbiome analysis reared at 35°C was identified as being positively infected by wet mount. Only statistically significant relationships are shown. A “*” indicates statistical significance below the “0.05” level.

Across time points, fish reared at 28°C exhibited the highest mean infection burden (4.86 worms per fish), followed by those at 32°C (3.6 worms per fish). Notably, at 14 dpe, fish at 32°C had a slightly higher infection burden (3.3 worms per fish) than those at 28°C (2.3 worms per fish). Tukey’s post hoc test revealed that infection burden was significantly higher in fish reared at 28°C and 32°C compared to those at 35°C (*P* < 0.05; Fig. 3B; Table S3B.3). Only a single larval worm was detected by wet mount in two fish from the 35°C group, while histological examination revealed a slightly higher prevalence, with larval worms observed in 9 out of 32 fish at this temperature (Fig. 3A; Table S3A.1). These results indicate that infection burden is highest at lower water temperatures. We also examined the development of mature female worms across temperature conditions. At 28°C, mature female worms were first detected at 28 dpe in 7 fish, whereas at 32°C, mature females were only observed in 4 fish. Interestingly, at 14 dpe, a single mature female was identified in a fish reared at 32°C, marking the earliest recorded instance of worm maturation at this temperature.

Additionally, we compared the sensitivity of infection detection between histology and wet mount methods on a subset of fish selected for microbiome analysis (*n* = 120; Fig. S3C; Table S3C.1). McNemar’s test revealed significant differences in detection sensitivity under specific conditions. At 35°C and 21 dpe, histology identified significantly more infections than wet mount (χ² = 4.17, *P* < 0.05; Fig. S3C; Table S3C.3), with 6 samples testing positive by histology alone compared to 0 by wet mount alone. No statistically significant differences were observed at other temperature and dpe combinations (*P* > 0.05; Table S3C.3). In cases where all samples were concordant (e.g., 28°C at 28 dpe and 35°C at 28 dpe), McNemar’s test could not be computed due to the absence of discordant pairs. These findings suggest that histological methods may be more sensitive than wet mounts, particularly at higher temperatures and intermediate time points. Collectively, these results suggest that higher water temperatures may have a protective effect against infection burden, limiting worm establishment and development in zebrafish.

### Gut microbiome response to parasite exposure varies across water temperature

To investigate how parasite exposure affects the gut microbiome under varying water temperatures, we analyzed fecal samples from exposed and control fish at multiple time points. *P. tomentosa* is known to alter the zebrafish gut microbiome (15), but it remains unclear how increasing water temperatures affect this response. We collected fecal samples for microbiome analysis of fish in the parasite exposed cohort at 14-, 21-, 28-, and 42 dpe. Similar to our parasite unexposed fish microbiome analyses, we built generalized linear models (GLM) to determine if temperature, time or their combination associated with variation in measures of microbial diversity and composition of parasite exposed fish (Table S4A.1). An ANOVA test of these GLMs revealed that alpha-diversity varied as a function of temperature for all measures (P<0.05; Fig. 4A; Table S4A.2). A post hoc Tukey test clarified that gut microbial diversity between 28°C and 32°C water temperature reared fish significantly differed for all alpha-diversity metrics (P<0.05; Table S4A.3), and gut microbial diversity differed between 28°C and 35°C water temperature reared fish as measured by Simpson’s Index. However, we did not find significant differences in gut microbial diversity between 32°C and 35°C water temperature reared fish for all alpha-diversity metrics, or between 28°C and 35°C water temperature reared fish as measured by Shannon Entropy, richness and phylogenetic diversity metrics. These results indicate that moderate increases in water temperature promotes gut microbial diversification in parasite exposed fish, but diversification of gut microbes plateaus in parasite exposed fish reared at higher water temperatures.

**Figure 4.**
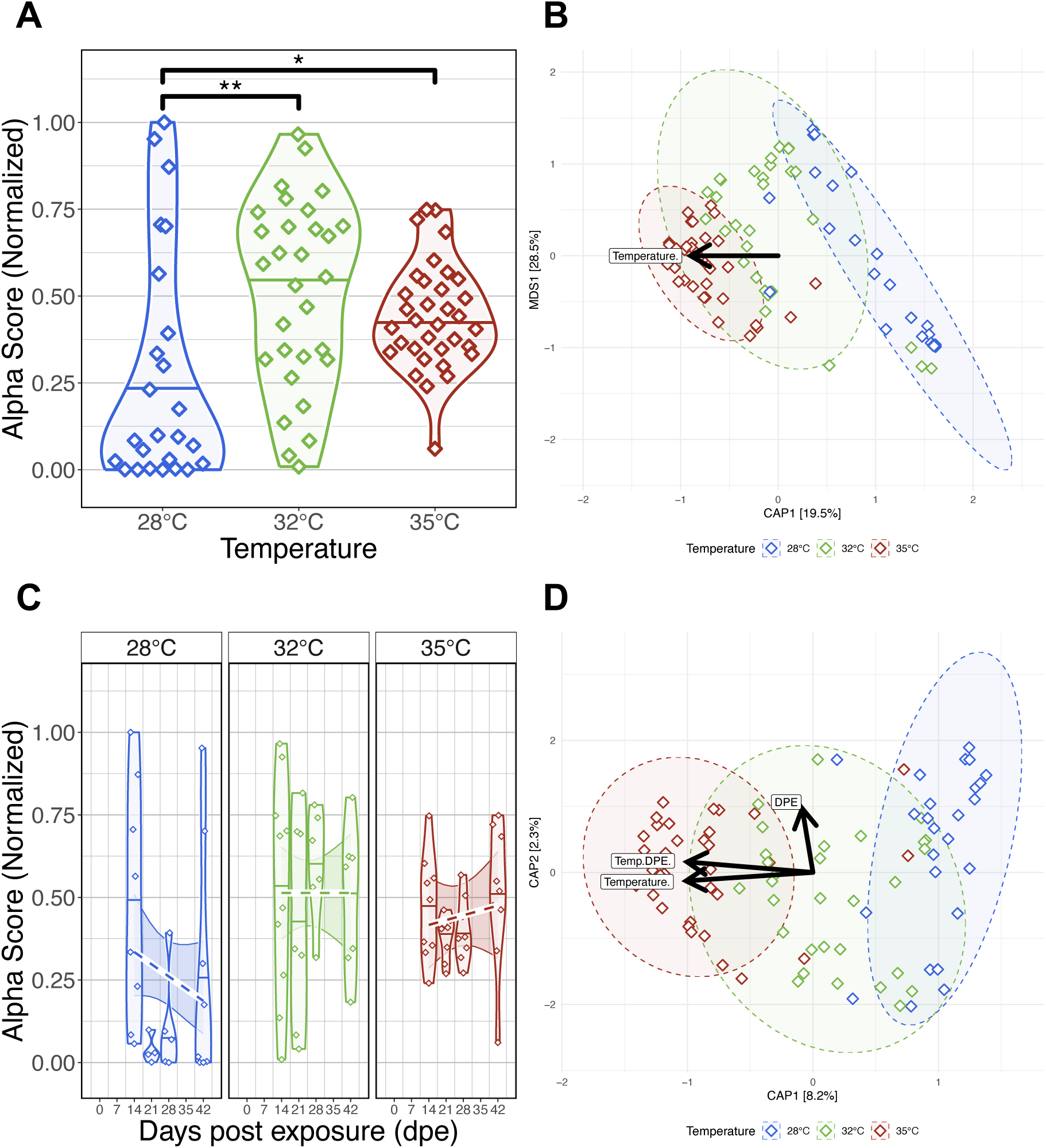
Effects of *Pseudocapillaria tomentosa* exposure on zebrafish gut microbiomes reared at different water temperatures. (A) Simpson’s Index of diversity shows that gut microbiome diversity significantly differs between fish reared at 28°C water temperature to fish reared at 32°C and 35°C water temperatures. (B) Capscale ordination based on the Bray-Curtis dissimilarity of gut microbiome composition constrained on the main effect of temperature. The analysis shows that gut microbiome composition significantly differs between parasite exposed fish reared at different water temperatures. (C) Simpson’s Index of diversity shows microbial gut diversity decreases with time from 0 days post exposure (dpe) to 42 dpe in parasite exposed fish reared at 28°C water temperature. (D) Capscale ordination of gut microbiome composition based on the Canberra dissimilarity constrained on the main effects of water temperature and time (days post exposure, dpe), and their interaction. The analysis shows that shows that gut microbiome composition differs between parasite exposed fish across time depending on water temperature. Ribbons and ellipses indicate 95% confidence interval. Only statistically significant relationships are shown. A “*” indicates statistical significance below the “0.05” level. Black arrows indicate direction of greatest change in the indicated covariates.

For each beta-diversity metrics we considered, PERMANOVA tests found that temperature significantly explained the variation in microbiome composition in parasite exposed fish (PERMANOVA, P<0.05; Fig. 4B; Table S4B.1). These results indicate that gut microbiome communities of parasite exposed fish reared at the same water temperature are more similar to each other than fish reared at different water temperatures. Additionally, we found beta-dispersion levels were significantly elevated between water temperature groups (P<0.05; Table S4B.2). A post hoc Tukey test clarified that beta-diversity dispersion levels were highest in fish reared at 28°C, followed by fish reared at 32°C and 35°C water temperatures (P<0.05; Table S4B.3). These results indicate that that gut microbiome communities of parasite exposed fish reared at lower water temperatures are more inconsistent in composition than parasite exposed fish reared at higher water temperatures.

Next, we compared our results across five time points to evaluate how parasite exposure and water temperature impacted gut microbiome diversity and composition. An ANOVA test did not find significant main effects of time as measured by Shannon Entropy and Simpson’s Index (P>0.05; Table S4C.2), but found marginally significant main effects of time as measured by richness and phylogenetic diversity (P=0.064 and P=0.078; Table S4C.2). Moreover, linear regression did not reveal significant interaction effects between temperature and time across all alpha-diversity metrics (P>0.05; Fig. 4C; Table S4C.2). These results indicate increasing water temperatures generally do not consistently impact microbial gut diversification over time in parasite exposed fish, and particular microbial clades appear more sensitive to the effects of time depending on temperature.

PERMANOVA tests found that community composition was best explained by the interaction effects between temperature and time using the Canberra beta-diversity metric (PERMANOVA, P<0.05; Fig. 4D; Table S4D.1), but a significant interaction effect was not observed using the Bray-Curtis and half-weighted UniFrac dissimilarity metrics (P>0.05; Table S4D.1). Given how these metrics weigh the importance of rarer (e.g., Canberra) versus abundant (e.g., Bray-Curtis) microbial community members, these results indicate that abundant members of the microbiome community are more robust to the effects of temperature across time in parasite exposed fish, while rarer taxa are more sensitive to the effects of time depending on temperature. Moreover, a pairwise analysis of beta-dispersion found significantly elevated levels of dispersion between fish reared across different temperatures and time as measured by all beta-diversity metrics (P<0.05; Table S4D.2). These results indicate that parasite exposure inconsistently impacts gut microbial community composition across time depending on temperature (P<0.05; Table S4D.2). Collectively, these results indicate that parasite exposure can impact gut microbiome diversity and composition, and these impacts are greatest at lower temperatures.

### Gut microbiome response has a non-linear relationship with infection burden

Given the differences we observed in gut microbiome diversity and composition across parasite exposed fish reared at different water temperatures, we further investigated whether gut microbiomes of parasite exposed fish vary depending on presence of infection and infection burden. Linear regression did not find significant main effects of presence of infection or significant interaction effects between presence of infection and temperature on gut microbial alpha-diversity for all metrics we measured (P>0.05; Fig 5A; Table S5A.1-2). These results indicate that gut microbial diversity does not differ as a function of presence of infection. Moreover, a PERMANOVA analysis found microbial community composition was best explained by presence of infection as measured by Canberra (PERMANOVA, P<0.05; Fig. 5B; Table S5B.1), but a significant result was not observed by the other beta-diversity metrics we measured. Additionally, we did not find statistically significant results between the interaction effects of water temperature and presence of infection on gut microbial community composition. These results indicate that rarer members of microbial communities of parasite exposed fish vary by presence or absence of infection, but abundant microbes do not. However, we did detect elevated levels of beta-dispersion across fish reared at different water temperatures depending on presence of infection (P<0.05; Table S5B.2). These results indicate that gut microbiome composition inconsistently varies between fish depending on presence of infection and water temperature.

**Figure 5.**
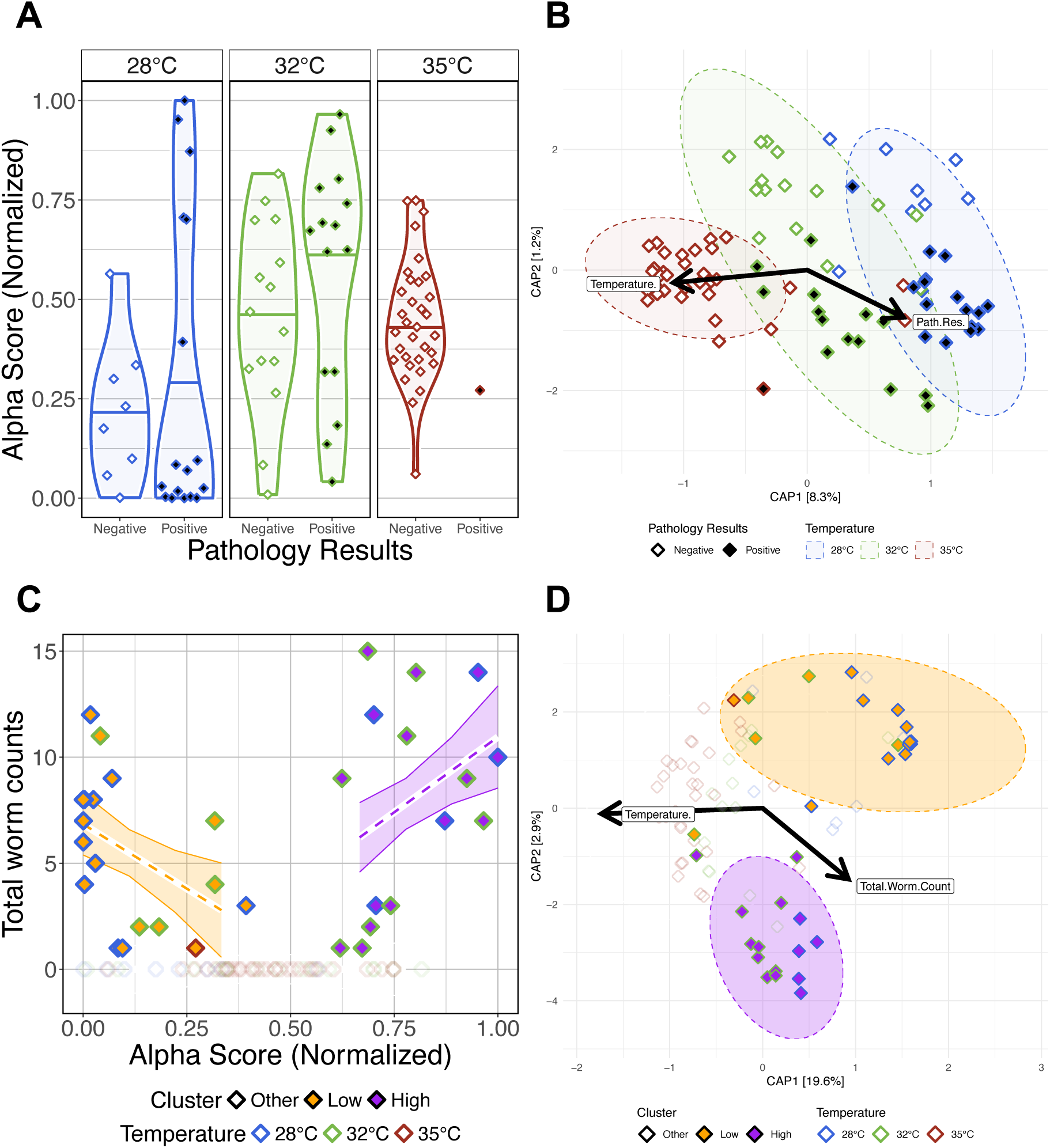
The impacts of presence of infection and infection burden on the gut microbiomes of *Pseudocapillaria tomentosa* exposed zebrafish. (A) Simpson’s Index for diversity of parasite exposed fish. Gut microbial alpha-diversity does not significantly differ between fish reared at the same water temperature depending on presence of infection. (B) Capscale ordination based on the Canberra dissimilarity of gut microbiome composition of parasite exposed fish constrained on the main effects of temperature and pathology result. The analysis shows that gut microbiome composition significantly differs between positively infected fish reared at different water temperatures. (C) Infection burden (total worm counts) is positively correlated with lowest or highest alpha diversity scores in positively infected fish. (D) Capscale ordination based on the Bray-Curtis dissimilarity of gut microbiome composition constrained on the main effects of water temperature and infection burden. The analysis shows that gut microbiome composition significantly differs between clusters of Low, High and Other fish. Samples points are colored by water temperature, and filled by “Cluster” grouping. Samples with at least one detectable worm and an alpha-diversity score less than 0.5 are categorized as Low (orange fill), samples with at least one detectable worm and an alpha-diversity score greater than 0.5 are categorized as High (purple fill), and samples with no observable infection are categorized as Other (white and transparent fill). Ribbons and ellipses indicate 95% confidence interval. Only statistically significant relationships are shown. A “*” indicates statistical significance below the “0.05” level. Black arrows indicate statistically significant covariates and direction of greatest change in the indicated covariates.

Next, we investigated how infection burden (i.e., number of intestinal worms detected) impacted parasite exposed fish gut microbiome diversity and composition. We used GLMs to determine if infection burden associated with variation in gut microbial alpha-diversity (Table S5C.1). An ANOVA test of these GLMs revealed that alpha-diversity varied as a function of infection burden as measured by Shannon Entropy and Simpson’s Index (P<0.05; Table S5C.2.2), but the interaction effects between infection burden and water temperature did not significantly explain variation in alpha-diversity across all measures (P>0.05; Table S5C.2.2). These results indicate that gut microbial diversity varies as a function of parasitic worm count. A PERMANOVA analysis found microbial community composition was best explained by infection burden as measured by all beta-diversity metrics (PERMANOVA, P<0.05; Table S5C.2.1), but the interaction effect between infection burden and temperature was not significant (P>0.05; Table S5C.2.1). These results indicate that higher infection burden drives increased inconsistency in gut microbial composition, regardless of water temperature.

Upon closer inspection of our infection burden results, we observed a non-linear relationship between infection burden and alpha-diversity scores, where highest infection burden associated with either highest or lowest alpha-diversity scores. To further explore this non-linear relationship between gut microbial diversity and infection burden, we grouped parasite exposed fish samples based on their alpha-diversity scores and infection burden. Parasite exposed fish samples with at least one intestinal worm detected were classified as “Low” or “High” if their alpha-diversity score was less than or greater than the median alpha-diversity score, respectively. Fish with zero detected worms were classified as “Other”. When grouping samples either Low or High based on alpha-diversity scores as measured by the Simpson’s index, we find that the samples in the Low group are composed of 67% 28°C and 33% 32°C water temperature reared fish, samples in the High group are composed of 33% 28°C and 67% 32°C water temperature reared fish, and samples in the Other group are composed of 14% 28°C, 27% 32°C, and 59% 35°C water temperature reared fish (Table S5C1.0). These results indicate that group membership trends with water temperature. To test this supposition, we used GLMs to determine if infection burden associated with variation in alpha-diversity score grouping (Table S5C.1.1). An ANOVA test of these GLMs revealed significant main effects of group for all alpha-diversity measures (P<0.05; Fig 5C; Table S5C.2.1), and significant interaction effects between group and alpha-diversity score. Notably, fish in the Low group had a significant negative slope and fish in the High group had a significant positive slope between alpha-diversity and infection burden as measured by Shannon Entropy and Simpson’s Index. These results indicate that parasite exposed fish have diverging gut microbial alpha-diversity responses to high infection burden.

Additionally, we find that these groups of samples – based on high versus low alpha-diversity scores of parasite exposed fish – also formed two distinct clusters in beta-diversity space. A PERMANOVA analysis detected significant clustering between Low, High, and Other groups across each measure of beta-diversity (PERMANOVA, P<0.05; Fig. 5D; Table S5D.1.1). However, this effect was weakest when considering the Canberra metric. Furthermore, a pairwise analysis of beta-dispersion finds significantly elevated dispersion levels between group membership as measured by Canberra metric, but not the other beta-diversity metrics (Table S5D.2). Given that the Canberra metric gives rarer taxa greater importance in its beta-diversity calculations than the other metrics we evaluated, these results suggest there is more consistency in microbial composition among abundant taxa within samples that share Low or High group membership, but not among more rarer taxa. A post-hoc Tukey test also clarified that beta-dispersion levels are significantly different between fish in the High and Other groups compared to fish in the Low group as measured by the Canberra metric (Table S5D.3). Together, these results indicate that rarer members of the gut microbiome are less consistently represented across fish in the Low cluster group as compared to fish in the High and Other cluster groups. Collectively, these results indicate that the microbiome response of fish with heaviest infection burden diverge into two distinct trajectories, which may be influenced by water temperature.

### Parasite exposure exacerbates water temperature differences in gut microbiome structure

Next, we sought to determine whether the gut microbiomes of zebrafish exposed to the parasite *Pseudocapillaria tomentosa* respond differentially compared to parasite unexposed control fish across increasing water temperatures. Prior to the parasite exposure at 164 dpf (or 0 dpe), we collected fecal samples from both cohorts of control and parasite exposed fish. Following fecal sample collection, fish in the parasite exposure group were exposed to *P. tomentosa*. We collected subsequent fecal samples at 14-, 21-, 28- and 42 dpe. Fecal samples were then measured for gut microbial diversity and composition and compared between parasite unexposed and exposed fish. We built generalized linear models (GLM) to determine if parasite exposure as a function of water temperature associated with microbial diversity and composition measures (Table S6A.1). Within pre-exposed (i.e., 0 dpe) samples, we did not observe any significant associations between the interaction effect of parasite exposure and water temperature across any of the alpha-diversity measures (P>0.05; Fig. 6A; Table S6A.2). These results indicate that at 0 dpe prior to parasite exposure, gut microbial diversity measures of fish reared at the same water temperature are not different from one another. Furthermore, PERMANOVA tests revealed significant differences in microbiome composition between control and pre-exposed fish across all beta diversity metrics. Homogeneity of dispersion tests revealed a significant difference in group variability for Bray-Curtis (P<0.05; Fig. 6B; Table S6B.2), but not for Canberra or Generalized UniFrac. Post hoc Tukey tests indicated no significant pairwise differences in dispersion for any metric (Table S6B.3), suggesting that the observed dispersion effect in Bray-Curtis was not driven by specific group outliers. Given that temperature alone consistently explained the variation in microbial communities followed by treatment effects, these results suggest that prior to parasite exposure microbial communities vary by water temperature and possibly stochasticity of tank effects.

**Figure 6.**
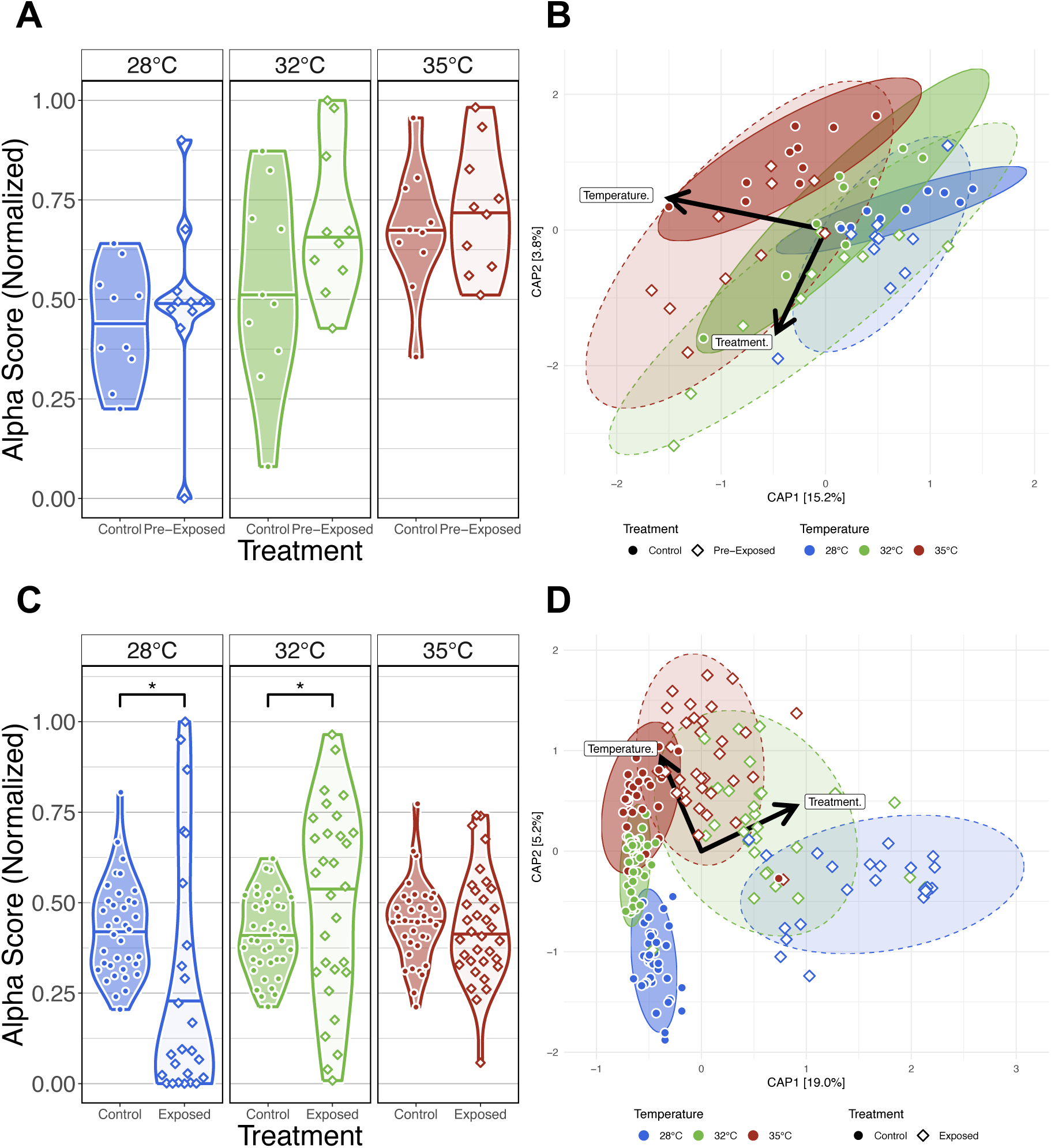
Comparison of the effects of water temperature on the gut microbiome between parasite exposed fish and parasite unexposed fish. (A) Simpson’s Index for diversity of parasite unexposed and pre-exposed fish at 0 days post exposure (dpe). Prior to parasite exposure gut microbial alpha-diversity does not significantly differ between fish reared at the same water temperature. (B) Capscale ordinations based on the Bray-Curtis dissimilarity of gut microbiome composition constrained on the main and interaction effects of temperature and parasite exposure (treatment) of pre-exposure samples at 0 dpe. (C) Simpson’s Index for diversity of parasite unexposed and exposed fish. Gut microbial alpha-diversity significantly differs between parasite exposed fish reared at 28°C and 32°C water temperature relative to unexposed control fish, but gut microbial alpha-diversity does not differ between parasite unexposed and exposed fish reared at 35°C water temperature. (D) Capscale ordinations based on the Bray-Curtis dissimilarity of gut microbiome composition constrained on the main and interaction effects of temperature and parasite exposure (treatment) of post-exposure samples after 0 dpe. The analysis shows gut microbiome composition differs between fish reared at different water temperatures prior to parasite exposure, and parasite exposure further drives these temperature associated differences in microbiome community composition. Ribbons and ellipses indicate 95% confidence interval. Ribbons and ellipses indicate 95% confidence interval. Only statistically significant relationships are shown. A “*” indicates statistical significance below the “0.05” level. Black arrows indicate statistically significant covariates and direction of greatest change in the indicated covariates.

We next compared our results between control and exposed fish across each water temperature to determine how parasite exposure impacts gut microbiome diversity and composition. Linear regression revealed microbial gut alpha-diversity was significantly associated with the interaction effect between temperature and treatment for any alpha-diversity metric we assessed (P<0.05; Fig. 6C; Table S6C.1-2). A post hoc Tukey test clarified that microbiome diversity was significantly different between exposure groups of fish reared at 28°C water temperature as measured by Simpson’s Index (P<0.05; Table S6C.3), at 32°C water temperature as measured by all alpha-diversity metrics (P<0.05; Table S6C.3), and at 35°C as measured by richness and phylogenetic diversity (P<0.05; Table S6C.3). These results indicate that gut microbial diversity differs between unexposed and exposed fish depending on water temperature, and parasite exposure uniquely impacts particular microbial clades, rare and abundant taxa depending on water temperature. Additionally, PERMANOVA tests found that microbiome composition differed between control and exposed fish reared at all water temperatures as measured by all beta-diversity metrics (P<0.05; Table S6D.1). These results suggest that the gut microbiomes compositions between control and parasite exposed differed in microbiome community composition regardless of water temperature. Moreover, a pairwise analysis of beta-dispersion found elevated levels of dispersion across all beta-diversity metrics measured, and dispersion levels were highest among parasite exposed fish reared at lower water temperatures (P<0.05; Table S6D.2). These results suggest that gut microbiome community composition is less consistent between parasite unexposed and exposed fish reared at lower water temperatures. Collectively, these results demonstrate that water temperature dictates how exposure to parasites alters the temporal trajectory of the gut microbiome.

### Gut microbial relative abundance significantly associates with environmental conditions and stressors

Finally, to evaluate how gut microbial abundance is influenced by environmental conditions and stressors (e.g., worm infections), we quantified differential abundance using MaAsLin2. Our analysis revealed 277 unique taxa at the Genus taxonomic level with at least one significant associations between taxon abundance and a covariate (Q<0.05, Fig. 7; Table S7A.1). We observed several taxa were significantly associated with the effect of water temperature. Fish reared at 35°C water temperature were enriched for 37 taxa, and depleted of 54 taxa relative to fish reared at 28°C water temperature. Fish reared at 32°C were enriched for 42 taxa, and depleted of 47 taxa relative to fish reared at 28°C water temperature (Fig. 7). Notably, *Aeromonas and Pseudomonas* bacterial abundance significantly associated negatively and positively with the effects of water temperature, respectively. *Aeromonas and Pseudomonas* are common members of the zebrafish gut microbiome (16,17), and these genera’s bacterial abundance has previously been observed to associate with the effects of water temperature in zebrafish (13). These results indicate that gut microbes are differentially selected for across varying water temperatures and time. We also measured how taxon abundance change over time regardless of water temperature. Over the course of 42 days, fish were enriched for 73 taxa and depleted of 36 taxa (Fig. 7). Notably, *Bosea* and *Cloacibacterium* bacterial abundance were negatively associated with the effects of time. *Bosea* and *Cloacibacterium* are common members of the zebrafish gut microbiome (16–18), and were also previously identified as having negative associations with the effects of time in zebrafish (15). These results indicate that particular members of the gut microbiome associate with time irrespective of water temperature.

**Figure 7.**
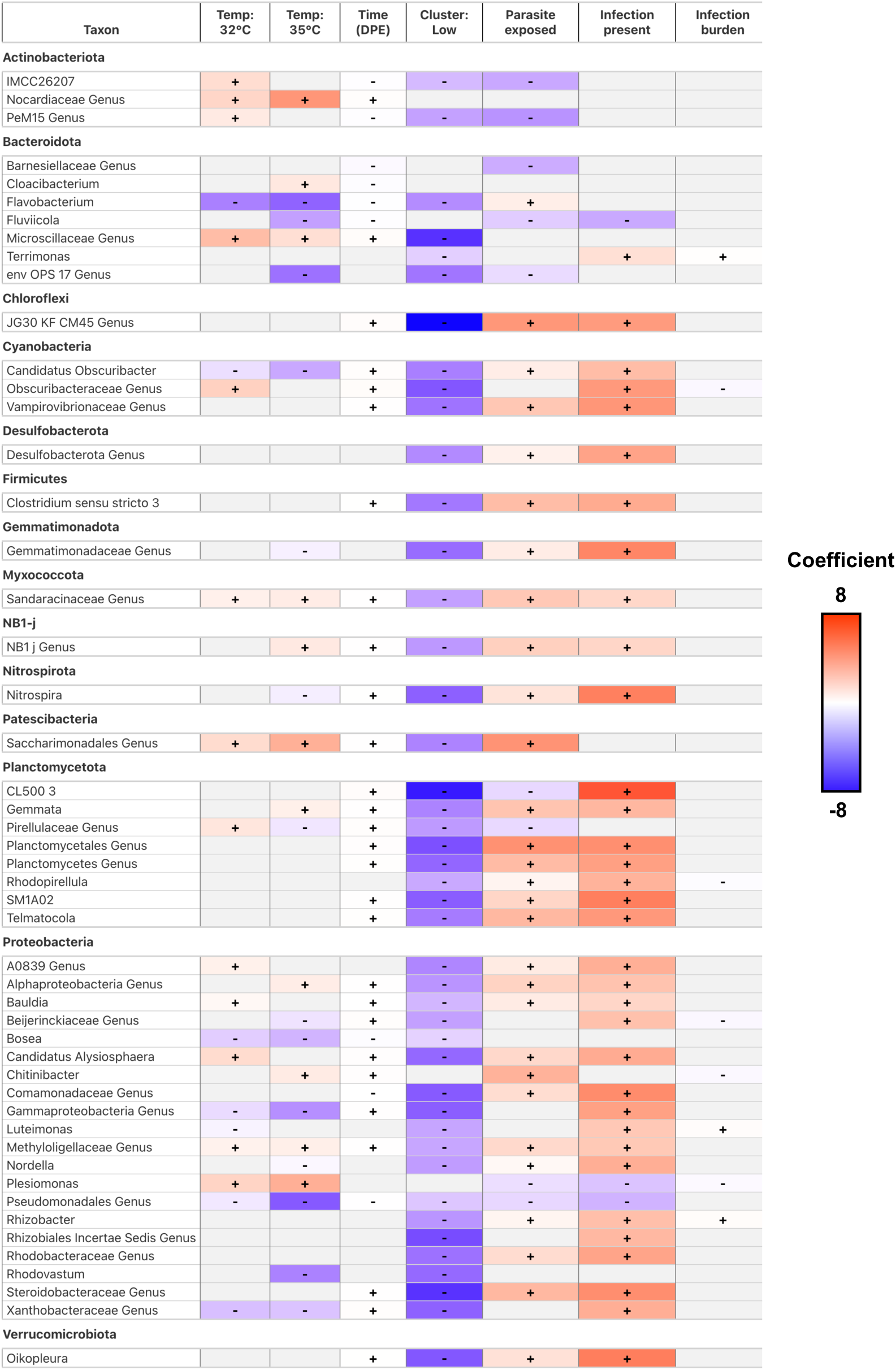
A heatmap of model coefficient values of the top 50 statistically significant abundant gut microbial taxa identified by MaAsLin2. The color of each cell represents the coefficient value and direction (red is positive, blue is negative). A “+” or “-” indicates a statistically significant association was observed between taxon abundance and a covariate. Gray colored cells indicate a significant effect was not observed.

Next, we sought to determine how parasite exposure impacted gut microbial abundance in fish. Fish exposed to *P. tomentosa* were significantly enriched for 74 taxa, and depleted of 35 taxa relative to parasite unexposed fish (Fig. 7). Notably, we find *Aeromonas*, *Chitinibacter*, and *Flavobacterium* are positively associated with parasite exposure, while *Plesiomonas, Phreatobacter* and *Cetobacterium* are negatively associated with parasite exposure. With the exception of *Phreatobacter* and *Cetobacterium*, these data are consistent with our prior work that found *P. tomentosa* exposure associated with altered bacterial abundance of these members of the zebrafish gut microbiome (15). We further investigated the effects of parasite exposure and measured how infection burden or presence of infection impacted gut microbial abundance. Fish with higher infection burden (i.e., number of parasitic worms present) enriched for 49 taxa and were depleted of 13 taxa, while fish that were positively infected enriched for 117 taxa and were depleted of 5 (Fig. 7). Notably, we find abundance of members of *Cetobacterium, Shewanella, Vibrio* and *Zooglogea* are negatively associated with infection burden. These taxa are identified as common members of the zebrafish gut microbiome (19). These results indicate that exposure to the intestinal parasitic worm *P. tomentosa* impacts common zebrafish gut microbiota, and presence of infection and severity have varying impacts to gut microbial abundance.

To deepen our analysis of parasite exposure on the zebrafish gut microbiome, we investigated how taxon relative abundance associated with gut microbiome diversity and composition. Previously, we found that parasite exposed fish reared at 28°C and 32°C water temperatures clustered into two distinct groups of community composition, which associated with high infection burden and either high or low alpha diversity scores. This observation led us to investigate which gut microbiota might be driving the clustering of the gut microbiomes of heavily infection burdened fish. We did not find significantly abundant taxa in the High group. We detected 1 taxon that was significantly enriched and 192 taxa that were significantly depleted among fish in the Low group (Fig. 7). Notably, we find *Aeromonas* was enriched, while *Mycobacterium* were depleted in the Low group fish. Some species of *Mycobacterium* are common pathogens in zebrafish facilities (20). These results indicate that the gut microbiome communities of parasite exposed fish experiencing heavy infection burden stratify into two distinct groups represented by the unique depletion of particular members of the gut microbiome. Collectively, these results indicate that environmental conditions associate with altered gut microbial abundance, and the response of specific members of the gut microbiome to environmental stressors varies depending on environmental conditions.

## Discussion

The zebrafish is an important model organism for understanding how environmental stressors impact the microbiome (17,18). Our work capitalized on the experimental control and scale afforded by the zebrafish model system to investigate how temperature and parasite exposure interact to influence infection and microbiome outcomes. While previous research has investigated how water temperature (13) and parasite exposure (15) independently impact the zebrafish gut microbiome, no studies in any *in vivo* experimental system, to our knowledge, have examined the microbiome’s temporal response to the combined effects of increasing water temperature and parasite exposure. Overall, we found that water temperature serves as a key contextual variable that dictates the severity of infection, the developmental state of worms, the composition of the gut microbiome in unexposed fish, and how the gut microbiome responds to parasite exposure and infection. These results underscore that the gut microbiome’s response to—and potentially its ability to buffer against—intestinal parasitic infection is influenced by other exogenous factors, in this case, water temperature. Furthermore, our findings challenge current expectations of climate change’s anticipated impact on aquatic organismal parasite burden (2,3). Consequently, it is important that we consider going forward how stacking multiple stressors, an experience inherent to life in the Anthropocene, may accelerate the arrival of dysbioses.

We found that parasitic infection burden was highest among zebrafish reared at ambient water temperatures. While field studies have documented arrested development in parasitic nematodes (e.g., *P. tomentosa*) during cold winter months (21), this study represents the first report of the effects of elevated temperature on parasitic nematode development in a fish host. Consistent with our prior work, temporal trends in *P. tomentosa* infection burden were similar for fish at ambient temperatures (15). However, contrary to expectations that elevated temperatures increase infection burden, we observed the opposite outcome: fish reared at the highest temperatures exhibited the lowest infection burden, with only a few larval-stage worms detected. Because parasite eggs were larvated at ambient temperature before being transferred to warmer tanks, we hypothesize that elevated temperatures may have impaired hatching once the eggs were ingested, reducing overall abundance of worms. Nevertheless, at 35°C, worms that did establish infections persisted but remained in an arrested state out to 28 and 42 days post-exposure (dpe), whereas at 28°C and 32°C, worms completed development and mated within 3 to 4 weeks, consistent with previous observations (14). These findings suggest that worm hatching and development may be temperature-sensitive, adding to prior research showing that many fish pathogens, from viruses to parasies, exhibit well-defined upper thermal boundaries for infectivity and pathogenesis. For example, the development is of the ciliate *Ichthyophthirius multifiliis,* a common cause of disease in aquarium fishes, is arrested by elevating water temperatures above 30°C (22). While we are not aware of this phenomenon with nematode parasites of fishes, analogous examples exist in terrestrial nematodes, such as *Wuchereria bancrofti*, where larval development is restricted to temperatures below 31°C in mosquitoes (23). Our findings address a gap in understanding of nematode development at a host’s upper thermal limit in an aquatic organism, and offers new insights into how climate change may disrupt aquatic disease ecological dynamics. Future research should investigate whether arrested development in *P. tomentosa* is a direct effect of temperature or mediated by host physiological responses.

Beyond direct effects on parasite development, poikilothermic (i.e., animals with variable body temperature and the inability to regulate it) hosts may gain protections against infection through temperature-dependent immune responses or gene expression changes. While studies on zebrafish immunity under elevated temperatures are limited, prior research in teleosts indicates that immune responses are host- and environment-specific, varying with the direction and duration of temperature shifts (24,25). For example, Dittmar et al. found that immune activity was highest at thermal limits and inversely related to acute temperature shifts in three-spine sticklebacks (26), whereas Bailey et al. observed suppressed immunity and increased parasite burden in rainbow trout exposed to chronic upper optimal thermal ranges (27). Although these studies differ in exposure regimes to ours, they highlight that colonization resistance may be influenced by temperature-sensitive immune responses and gene expression. Future research integrating immune function, gene expression, and histopathological assessments will be crucial to disentangling the host’s role in colonization resistance under chronic parasite exposure and elevated temperatures. Notably, controlled temperature manipulation is already used to mitigate certain aquaculture pathogens, such as *Ichthyophthirius multifiliis (22)*, where increasing tank temperature to 30°C can eliminate infections in susceptible fish. Our findings suggest that similar interventions may help mitigate or delay parasite infection in aquaculture settings.

We also found that zebrafish gut microbiome structure stratified depending on the environmental conditions of increasing water temperature. Our results are congruent with previous research that found increased water temperatures altered zebrafish gut microbial diversity and composition (13). Moreover, Wang *et al.* observed that zebrafish reared at different water temperatures manifested distinct liver carbohydrate metabolism profiles and temperature-dependent sensitivity to irradiation. A unique aspect of our study considered how the gut microbiome temporally varies as a function of water temperature. We found that water temperature acts as a filter on initial zebrafish gut microbiome assembly, and these initial differences in assembly between water temperature remained stable across time. Beyond zebrafish, analogous investigations have investigated how temperature variation shapes gut microbiome composition and function in mammals, fish, and other animal species (28,29). In particular, a recent meta-analysis of aquatic organisms’ response to temperature found similar, but inconsistent results to our study, wherein increasing water temperature is associated with both increases and decreases to gut microbial diversity, differences in gut microbiota community composition, and altered gut taxon abundance (29). Inconsistencies between prior work and ours could be driven by differences in magnitude of the stressor (i.e., press vs pulse; (30)), host species (31), facility or habitat effects (29,32,33), or diet (20). Despite these differences, the results of prior studies in conjunction with ours are consistent with the concept of environmental conditions acting as a habitat filter to shape initial gut microbiome assembly (34) and illicit environmentally dependent responses to exogenous stressors.

Finally, we observed a nonlinear relationship between gut microbiome diversity and infection outcomes, with water temperature moderating these dynamics. Consistent with our prior research on zebrafish infected by *Pseudocapillaria tomentosa* (15) exhibiting dysbiotic microbiomes described by the Anna Karenina Principle (AKP) (35). The AKP postulates that dysbiotic microbiome communities display higher rates of dispersion or are more inconsistent between dysbiotic individuals, whereas the microbiome community composition of healthy individuals are less dispersed or are more similar to one another. In particular, we observed elevated microbial community dispersion among heavily parasite infected fish, reflecting unstable microbial states characteristic of the AKP. However, when accounting for water temperature, the AKP effect diminished, and multiple alternative stable states emerged, which may explain inconsistencies in prior studies that failed to detect the AKP or found contrary results (29). Our findings underscore the importance of accounting for individuals’ temporal and spatial contexts to adequately assess microbiome stability in response to stressors (36). However, current definitions of microbiome stability (11,37,38), which rely on a homeostatic framework, may be insufficient to describe these dynamic shifts. A homeorhetic framework (39), conceptualizing stability as a change along a stable trajectory rather than a fixed state, could improve our ability to measure microbiome stability and dysbiosis across dynamic temporal and spatial scales. Shifting to such a model may reconcile discrepancies in AKP detection across studies and provide deeper insights into how microbiomes respond to exogenous stressors in different host systems.

In conclusion, we found that water temperature alters the contextual landscape of the microbiome to impact its response to an exogenous stressor of an intestinal parasite. Our work revealed that differences in environmental conditions of water temperature were sufficient to temporally change the gut microbiome’s response to parasitic exposure and impact infection outcomes in zebrafish. While the zebrafish gut microbiome differs taxonomically from other animal-microbiome systems, a considerable amount of functional capacity is shared between animals (40). Thus, zebrafish serve as a powerful model for investigating how environmental changes and stressor exposures influence microbiomes and host health. Our findings have important implications for microbiome research in the context of climate change, demonstrating that rising temperatures may have unexpected effects on gut microbiomes and infection outcomes. Future work should further clarify how gut microbiomes and host responses buffer against combined environmental stressors, ultimately shaping health outcomes in vertebrates.

## Methods

### Fish husbandry

5D strain zebrafish embryos were obtained from the Sinnhuber Aquatic Resource Center at Oregon State University, and reared in our vivarium at Nash Hall (Corvallis, OR, USA). The vivarium is a single pass flow through, using dechlorinated city water. Fish were then randomly divided into twelve 2.8 L tanks. The temperature was recorded daily and the ambient temperature ranged from 27 to 28°C. All other water conditions were monitored weekly, pH was maintained at 7.6, total ammonia was not detected, and conductivity ranged from 102 to 122. Light in the vivarium was provided for 14 hours/day. Fish were fed Gemma Micro 300 (Skretting; Fontaine-les-Vervins, France) at 1.5% body weight twice daily, except on weekends or during exposure to parasitic eggs. One plastic aquatic plant piece, approximately six inches in length, was added to each tank for enrichment. The use of zebrafish in this study was approved by the Institutional Animal Care and Use Committee (IACUC) at Oregon State University (permit number: 5151).

### Temperature exposure

Fish at 5 month-old were randomly divided into 12 9.5-L tanks (approximately 25 fish/tank). Each tank was outfitted with a 50W (28 °C treatment only) or 100W HG-802 Hygger titanium aquarium heater (Hygger, Shenzhen Mago Co., Ltd., Shenzhen City, Guangdong Province, China). Four of the twelve tanks were assigned to each of the temperature treatments: 28°C, 32°C, or 35°C. Two tanks for each temperature were held as pathogen negative controls and two tanks were exposed to *Pseudocapillaria tomentosa* as described below. Fish were acclimated to the prescribed temperature treatments by increasing the heater thermostat settings by 1°C every two days until the final prescribed temperature was achieved. Two temperature logging thermometers, one for the six pathogen negative control tanks and one for the six *P. tomentosa* exposed tanks, were rotated through the tanks every two days on weekdays to monitor temperature at each temperature treatment. The average range recorded for the water temperature treatments was +/- 1°C.

### Pseudocapillaria tomentosa exposure

Eggs were collected from a population of infected zebrafish that we constantly maintain in our laboratory as described by Martins *et al.* 2017 (41). Eggs were allowed to larvate for 6 days at 28°C, and fish were exposed at 25 larvated eggs/fish. Water flow was turned off for 36h to enhance exposures, while an airstone was provided to each tank to maintain adequate oxygen levels. This was a lower exposure dose than many of our previous studies (14). Therefore, we enhanced exposure adding 1 L of water from a stock tank holding infected fish twice a day during the 36 h hour post exposure period. This additional water supplement was created by siphoning water from the bottom of the exposed stock fish tank because the infectious stage is a larvated egg, which sinks in water.

### Infection assessment

Exposed and control fish were collected and examined for worm prevalence, abundance and state of development using wet mounts of whole intestines as described in Schuster *et al.* 2023 (42). After recording observations in wet mounts, the individual intestine was preserved in Dietrich’s solution and intestines of 95 fish were processed for histology prepared as described in Gaulke *et al.* 2019 (15). Here we focused on selected samples from fish from the 35°C group as very few worms were detected by wet mounts in this group. Two stepwise sections, 50 um apart, were obtained from each block to enhance the possibility of larval worms

### Fecal collection

Five fish from each tank were randomly selected for fecal sampling at 0 dpe (n=60; 5 samples/tank), prior to parasite exposure. Subsequent fecal sampling took place at 14- (n = 54), 21- (n = 48), 28- (n = 47), and 42 (n = 51) dpe to parasites. Fecal material was collected as previously described (20). In brief, fish were transferred to 1.4 L tanks (1 fish/tank) containing ∼ 0.4 L of fish water at least 30 min after the last feeding of the day. Fish were left to defecate overnight and all fecal material was collected from each tank the following morning in a 1.5ml microcentrifuge tube. Fecal samples were immediately spun at 10k rpm for 2 min, excess tank water was removed, and samples were snap frozen on dry ice and stored at -80 °C until processing. However, not all fish produced a fecal sample for a variety of reasons. For instance, experiments involving fish have expected mortality, and fish which died prematurely did not produce fecal samples. Additionally, infection conditions may have prevented infected fish from producing a fecal sample. Instances where fish failed to produce a fecal sample are noted in the metadata sheet.

### Microbial 16S rRNA library preparation and sequencing

Microbial DNA was extracted from zebrafish fecal samples and 16S rRNA gene sequence libraries were produced and analyzed following previously described methods (43). DNA was isolated from fecal samples using the DNeasy 96 PowerSoil Pro DNA kits (Qiagen, Hilden, Germany), in accordance with the manufacturer’s directions. In brief, samples were subjected to bead beating for 10 minutes using the Qiagen TissueLyser II, spun a max speed in the centrifuge, supernatant was process using 96 well columns, and DNA was eluted with 100µl Tris buffer. The V4 region of the 16S rRNA gene was PCR amplified using dual-index 16S primers and protocols (44). PCR was performed using 1 µl of purified DNA, 2µl of a 5µM mix of the forward and reverse dual-index primers, 5µl of Platinum II Hot-Start PCR Master Mix (Thermo Fisher), Carlsbad, CA), and 2µl water with the following conditions, 94°C, 3m; (94°C, 30s; 50°C, 30s; 68°C, 1m)x 35; 68°C 10m. PCR products were visualized on a 1.5% agarose gel and quantified on the BioTek Synergy H1 Hybrid Multi-Mode Plate Reader using the Quand-iT 1X dsDNA HS Assy kit (Thermo Fisher, Carlsbad, CA, USA). A 100ng aliquot of DNA was selected from each of the 300 samples, the pooled DNA was cleaned using the QIAGEN QIAquick PCR purification kit, and quantified using Qubit HS kit (Carlsbad, CA). The quality of the pooled library was verified on the Agilent TapeStation 4200. The prepared library was submitted to the Oregon State University Center for Quantitative Life Sciences (CQLS) for paired end 2x300 bp read sequencing on an Illumina MiSeq System

### Bioinformatic processing

All microbiome DNA sequence analyses and visualization were conducted in R (v 4.3.3)(45). Raw reads were filtered for quality, merged, and assigned using the DADA2 R package (v 1.26.0) as previously described (46). In brief, forward and reverse reads were trimmed at 250 and 225 bp, respectively, subsequently merged into contigs, and subject to amplicon sequence variant (ASV) identification. ASVs unannotated at the Phylum level or identified as non-bacterial were removed, which resulted in 674 remaining detected ASVs. Samples containing reads below the minimum required read count (<5000) were dropped from downstream analysis. The final sample number for microbiome analysis was 260. Phylogenetic analysis was conducted using MOTHUR (v 1.46.1)(47) with default parameters as previously described (17). Phylogeny was inferred using FastTree2 (48), an approximately-maximum-likelihood method. Microbiome and sample data were contained in a Phyloseq object using the Phyloseq R package (49), and the tidyverse (v 2.0.0)(50) and microViz (v 0.12.1) R packages were used for downstream data processing, analyzing, and visualization (51). Code for bioinformatic processing are available at https://github.com/sielerjm/Sieler2025_ZF_Temperature_Parasite/.

### Microbiome diversity metrics

All microbiome analyses were conducted at the genera level unless otherwise noted. We estimated four alpha-diversity metrics for each microbiome fecal sample: Simpson (52), Shannon (53), phylogenetic diversity (Faith’s PD (54); ASVs), and richness. We also estimated beta-diversity between each pair of microbiome fecal samples using three metrics. These included Bray-Curtis (55), Canberra (56), and half-weighted generalized UniFrac (57).

### Statistical Analyses

All statistical analyses were conducted in R (v 4.3.3)(45) with a significance level of α = 0.05, and randomization procedures employed a fixed seed (42) to ensure reproducibility. Code for statistical analyses are available at https://github.com/sielerjm/Sieler2025_ZF_Temperature_Parasite/.

Using methods previously described, we assessed normality of alpha-diversity scores using Shapiro-Wilk test (45,58,59), transformed non-normal scores using Tukey’s Ladder of Powers (59,60) and normalized from 0 to 1 (20) before incorporation into linear models. We used generalized linear models (GLM), we assessed the relationship between alpha-diversity score and experimental parameters. Post hoc Tukey Tests evaluated pairwise comparisons of models using the multcomp R package (v 1.4-25)(61). We corrected for multiple tests using Benjamini-Hochberg correction (62). Two-way ANOVA was used to determine if the expanded models of these GLMs significantly improved the response variable relative to the null model.

Beta-diversity models were generated using methods described previously (20). In brief, we assessed the relationship between experimental parameters and beta-diversity by applying a step-wise model selection approach as implemented in the capscale function (vegan R package v 2.6-4)(63). Optimal models were subsequently subject to permutation analysis of variance (PERMANOVA) with anova.cca using the vegan R package to determine if the selected model parameters significantly explained the variation in microbiome composition across samples. Differential abundance was measured using MaAsLin2 (64). We assessed beta-diversity dispersion within groups with betadisper using the vegan R package.

To assess the relationship between parasite infection outcomes and experimental parameters, we used negative binomial generalized linear models (GLM) with the glm.nb function from the MASS R package (v 7.3-60.0.1)(65) and used the negative binomial distribution to account for overdispersion in the count data, a common characteristic of parasite infection data (14). Significance of main effects and interactions was assessed using two-way ANOVA implemented through the Anova function with the Car R package (v 3.1-2)(66). Post-hoc comparisons were conducted using Tukey’s HSD tests via the emmeans package, where we estimated marginal means and performed pairwise contrasts with p-value adjustment using the Tukey method (67). Detection method comparisons were analyzed on a subset of samples used for microbiome analysis (120 samples, 20 samples/tank, ∼10 samples/time point; Table S3C.1). To compare detection methods between wet mount and histology, we used McNemar’s test (68), with discordant pairs (wet only vs histology only) examined at each temperature and DPE combination through 2×2 contingency tables Table S3C.2-3). Using similar methods as described above, we assessed the relationship between infection outcomes and microbiome diversity using GLMs (Table S3B.1).

## Abbreviations

dpe: days post exposure

## Acknowledgements

The authors thank the members of the Oregon State University Center for Quantitative Life Sciences for technical assistance with sequencing and maintenance of our computational infrastructure, and Dr. Corbin Schuster and Kelan Elliot for sample collection assistance.

## Data Availability

All code generated during this analysis is available in the GitHub repository at the following URL: https://github.com/sielerjm/Sieler2025_ZF_Temperature_Parasite. Supplementary tables and figures can be in the accompanying supplementary files. The raw sequence files generated during the current study are available at the NCBI Sequence Read Archive (SRA) project number: PRJNA1219243.

## Author contributions

TJS and MLK conceived and designed the study. CEA and MJS conducted the experiments. MJS, TJS, and KDK performed the gut microbiome and integrated analyses. MJS, TJS, MLK, CEA, KDK, contributed to the preparation and editing of the manuscript. MJS prepared the figures. All authors read and approved the final manuscript.

## Declarations

The authors declare no competing interests.

