## Supplemental Tables and Figures for "Modeling the zebrafish gut microbiome’s resistance and sensitivity to climate change and parasite infection": Supplemental_Figures__BioRxiv_FINAL.pdf

***7) Gut microbial abundance is significantly associated with environmental conditions and stressors* .................................................................................................. *Error! Bookmark not defined.***

### 2) Water temperature shapes gut microbiome structure

## S2A)

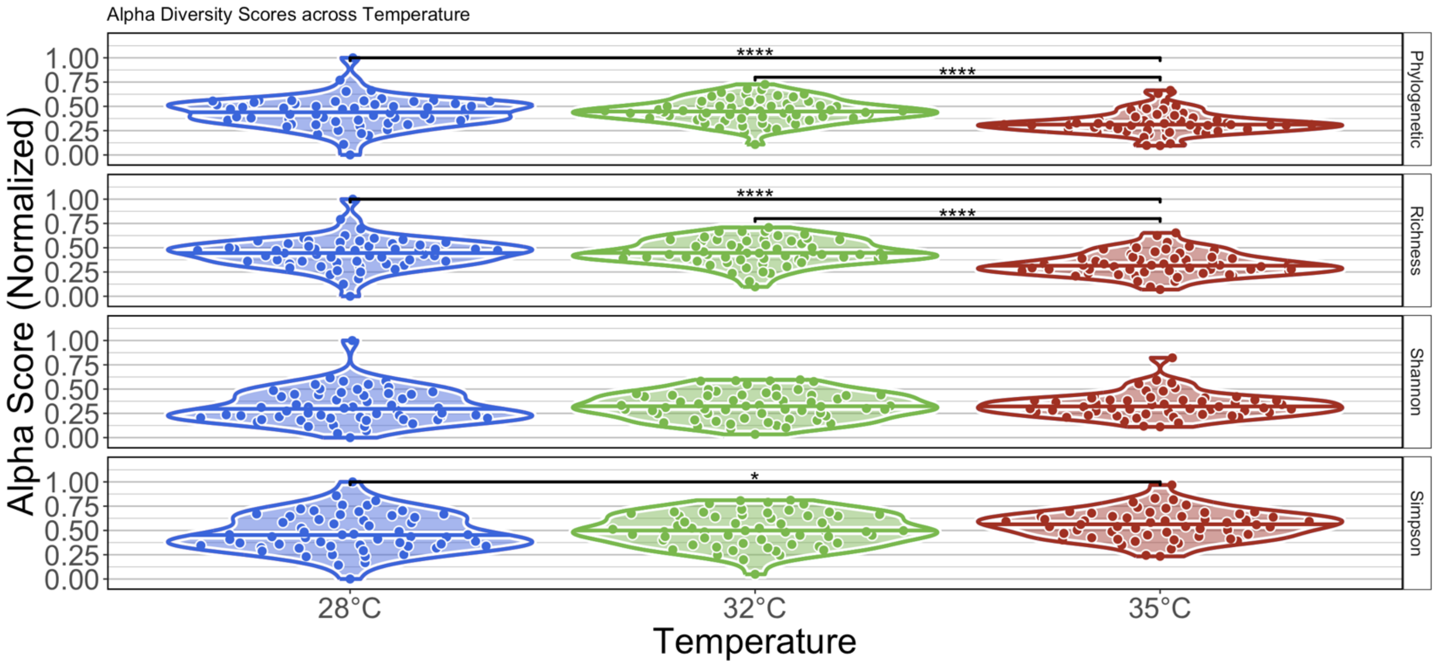

## S2B)

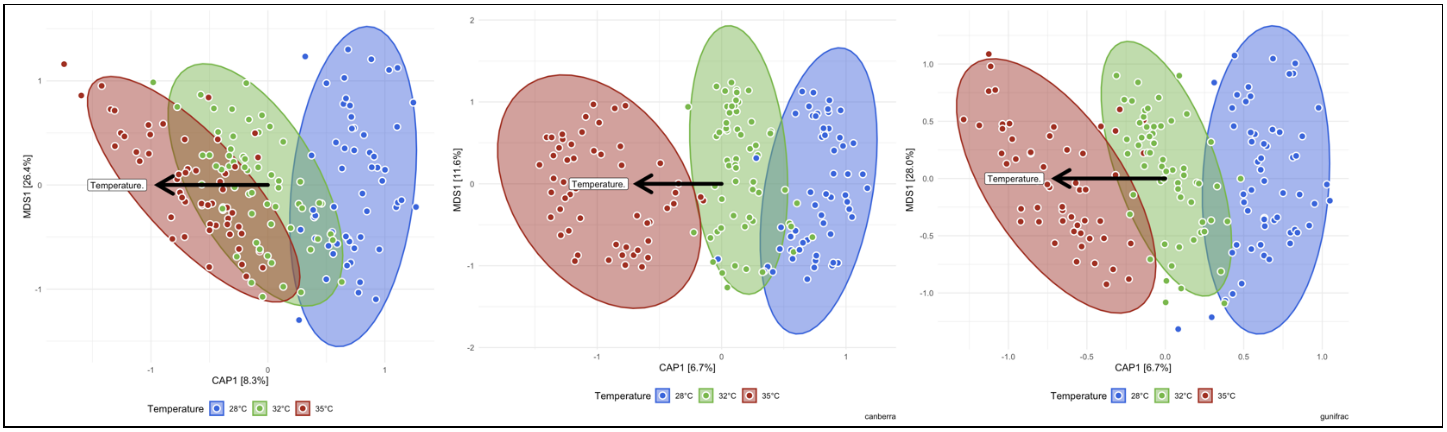

## S2C)

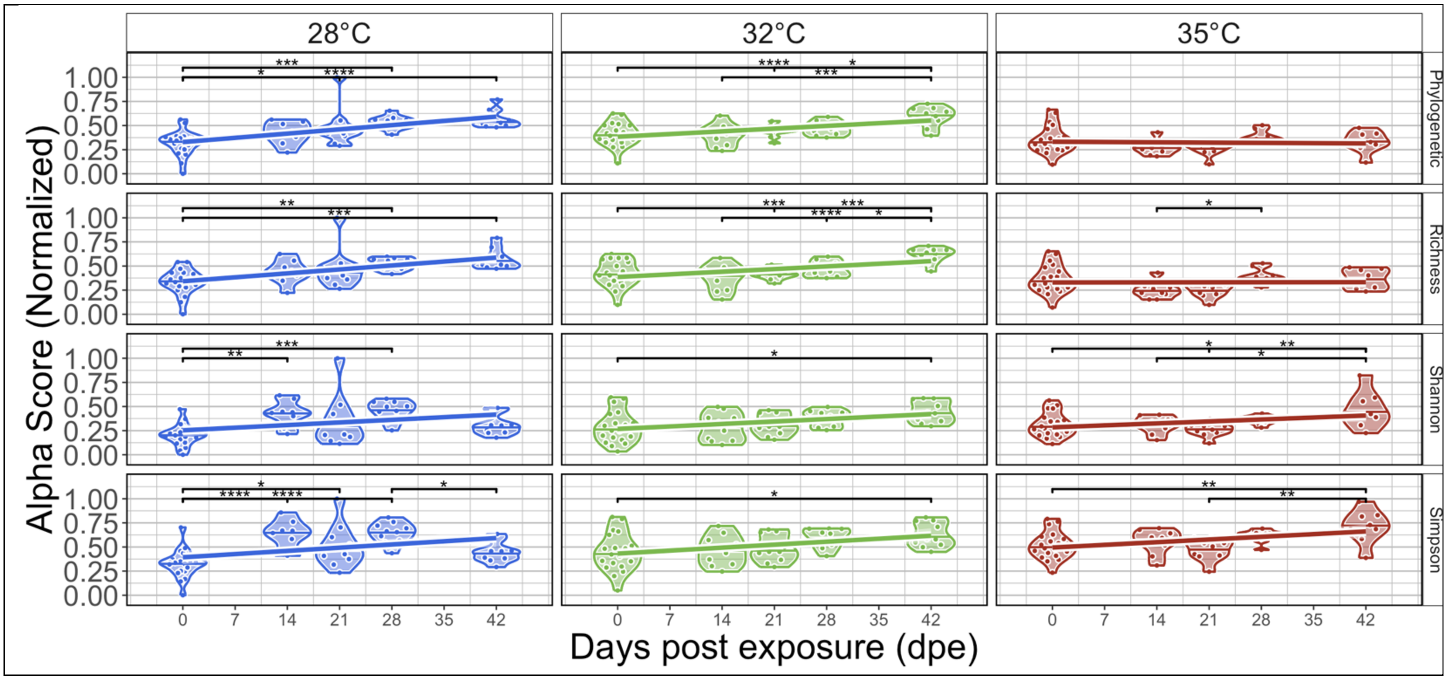
28°C
32°C
35°C
Phylogenetic
Richness
Shannon
Simpson
Alpha Score (Normalized)
Days post exposure (dpe)

S2D)

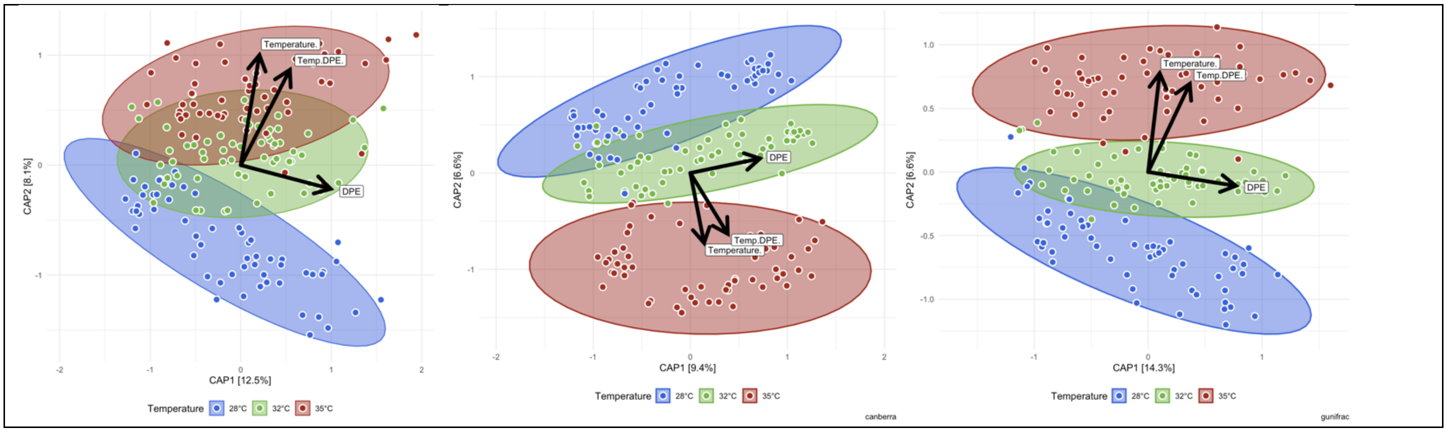
Temperature.
Temp.DPE.
DPE
CAP1 [12.5%]
CAP2 [8.1%]
CAP1 [9.4%]
CAP2 [6.6%]
CAP1 [14.3%]
CAP2 [6.6%]
Temperature
28°C
32°C
35°C
canberra
gunifrac

#### 3) Infection burden is highest in fish reared at lower water temperatures

S3A)

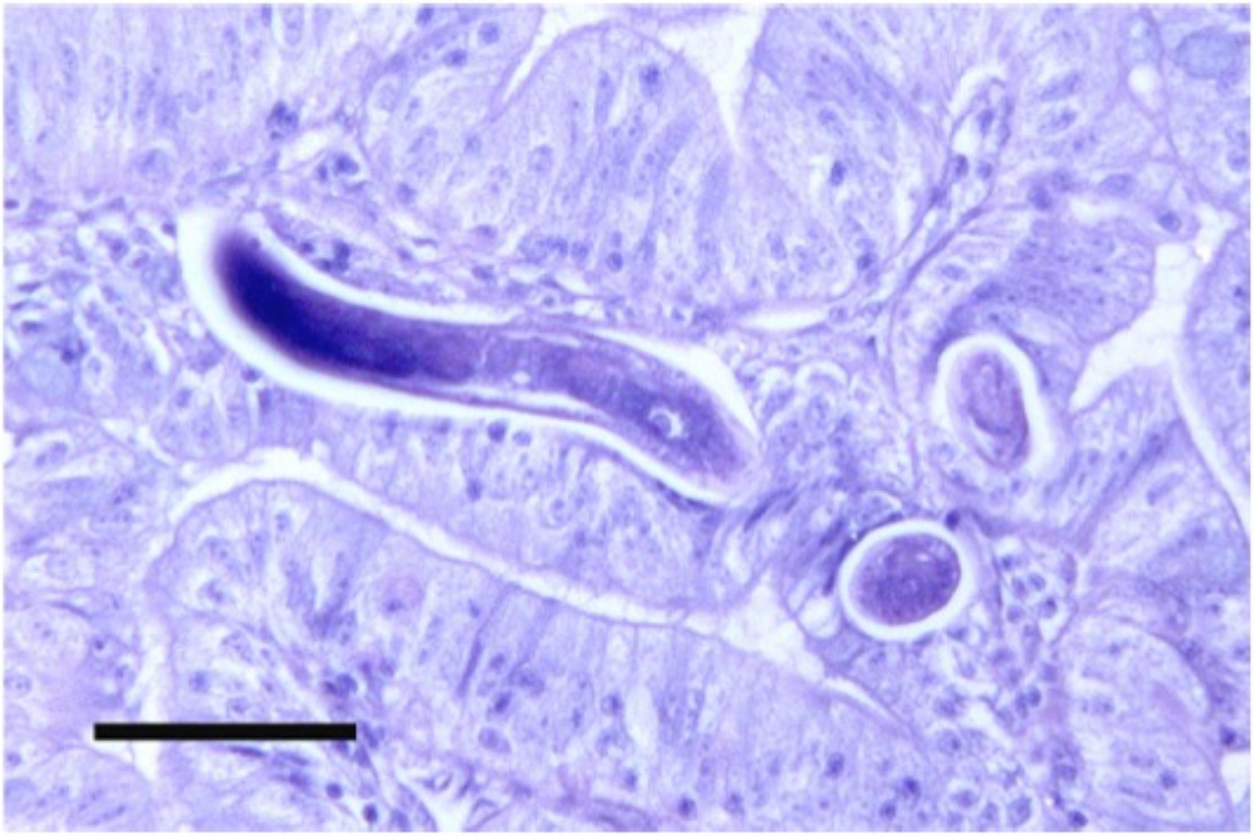

## S3B)

### S3B.1)

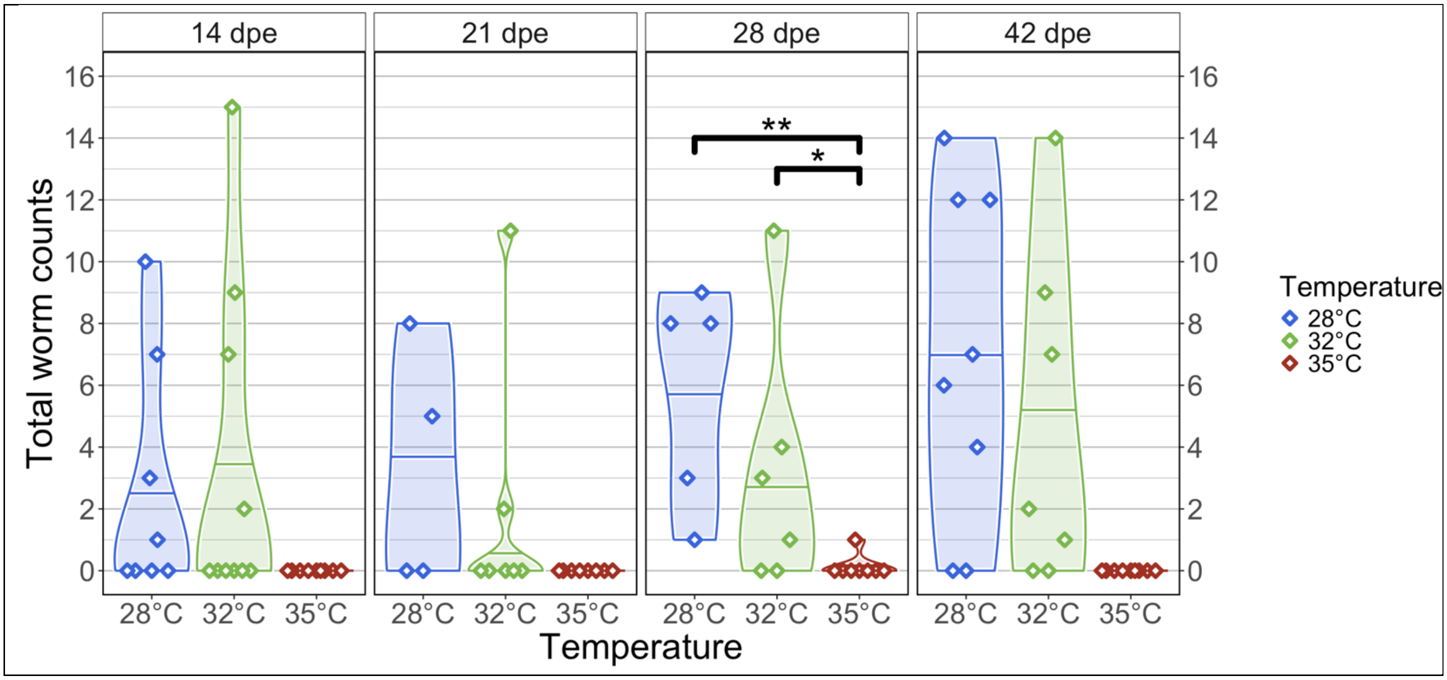

### S3B.2)

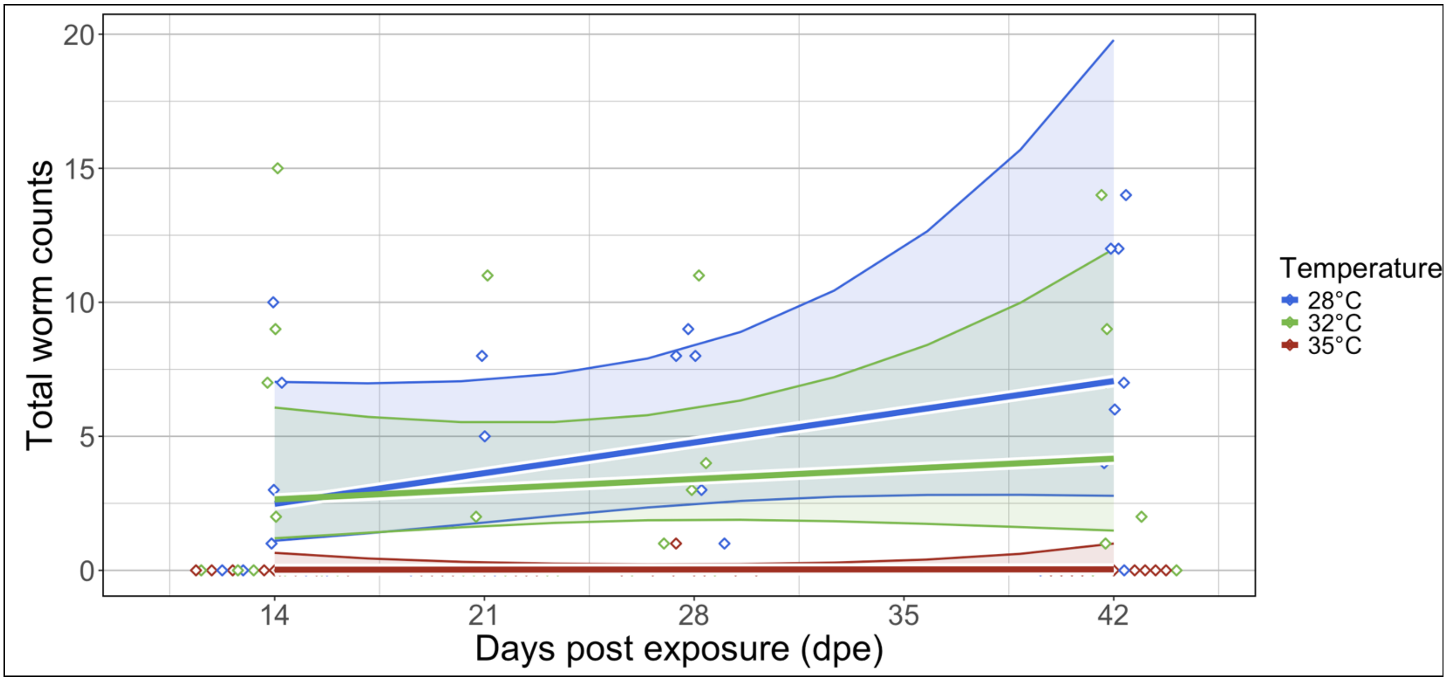

S3C)

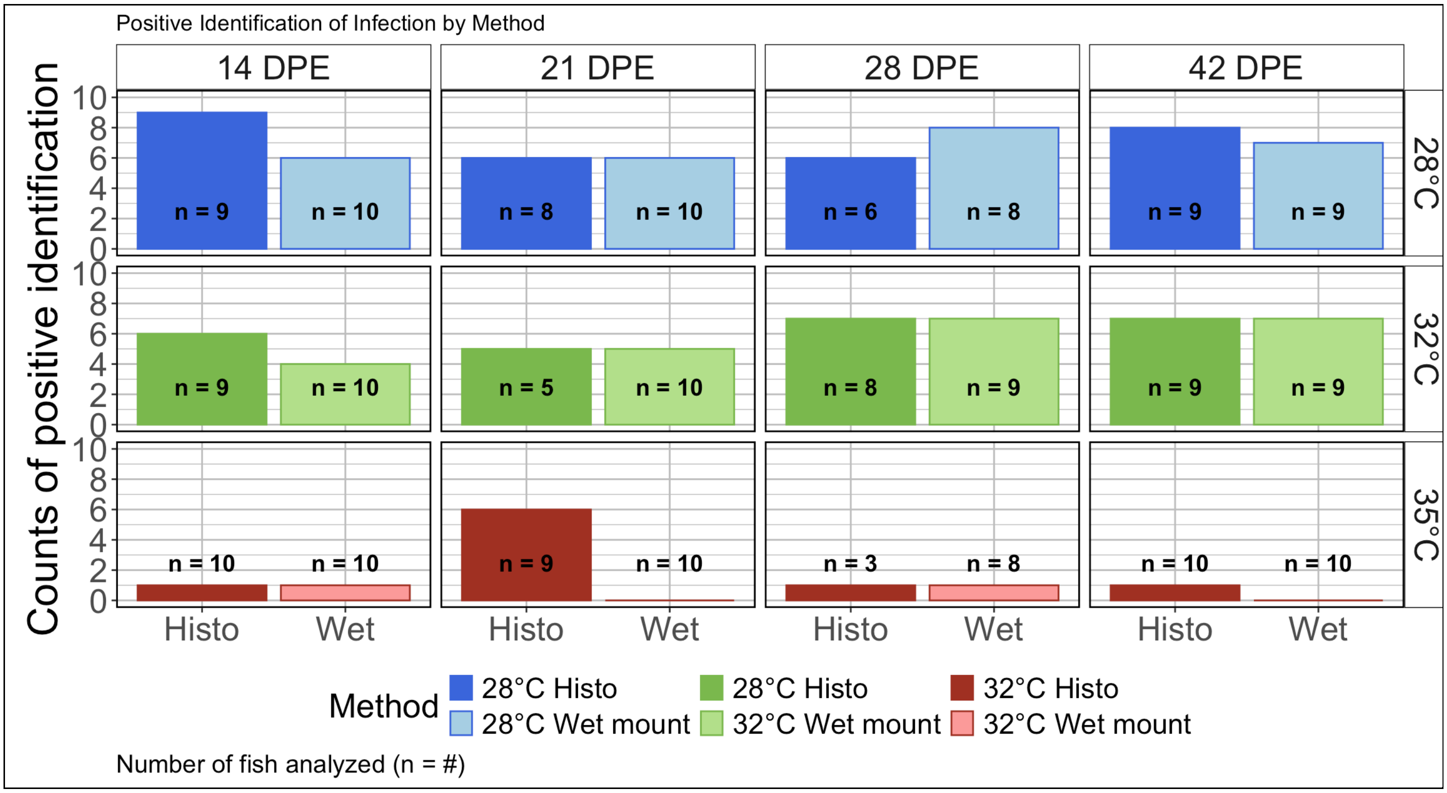

Positive Identification of Infection by Method
14 DPE
21 DPE
28 DPE
42 DPE
28°C
32°C
35°C
Counts of positive identification
0
2
4
6
8
10
n = 9
n = 10
n = 8
n = 10
n = 6
n = 8
n = 9
n = 9
n = 9
n = 10
n = 5
n = 10
n = 8
n = 9
n = 9
n = 9
n = 10
n = 10
n = 9
n = 10
n = 3
n = 8
n = 10
n = 10
Histo
Wet
Method
28°C Histo
28°C Wet mount
28°C Histo
32°C Wet mount
32°C Histo
32°C Wet mount
Number of fish analyzed (n = #)

#### 4) Gut microbiome response to parasite exposure varies across water temperature

### S4A)

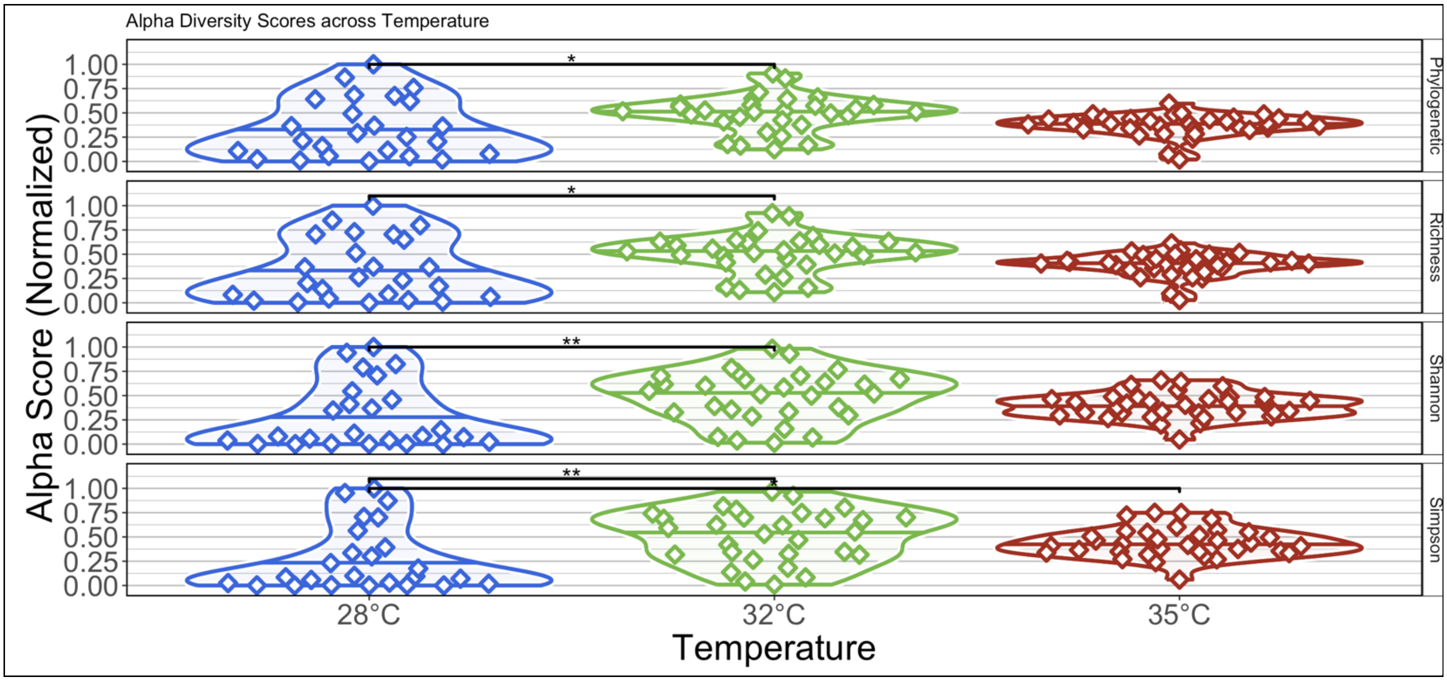

### S4B)

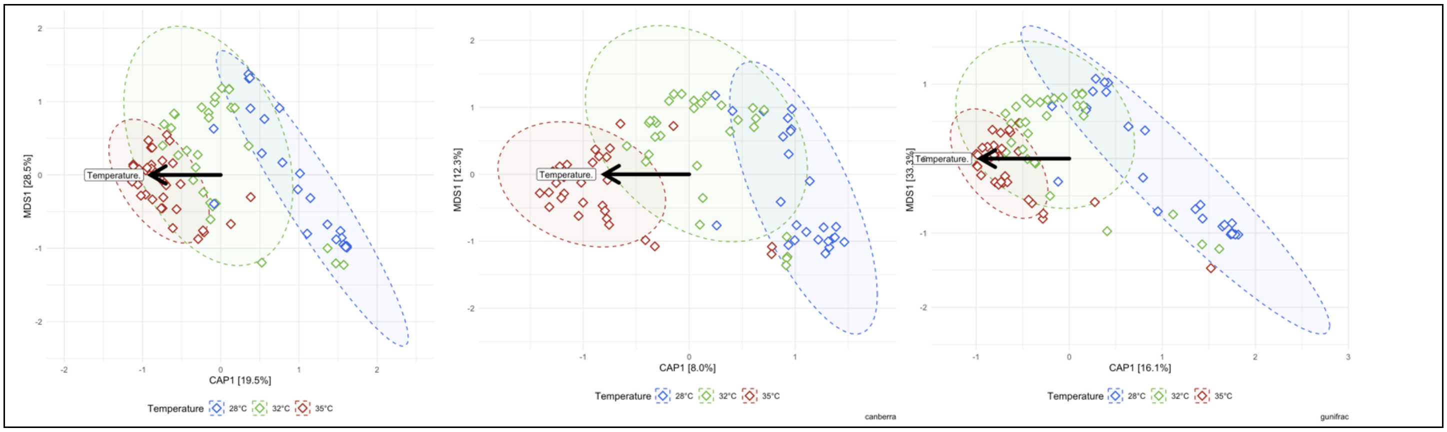

S4C)

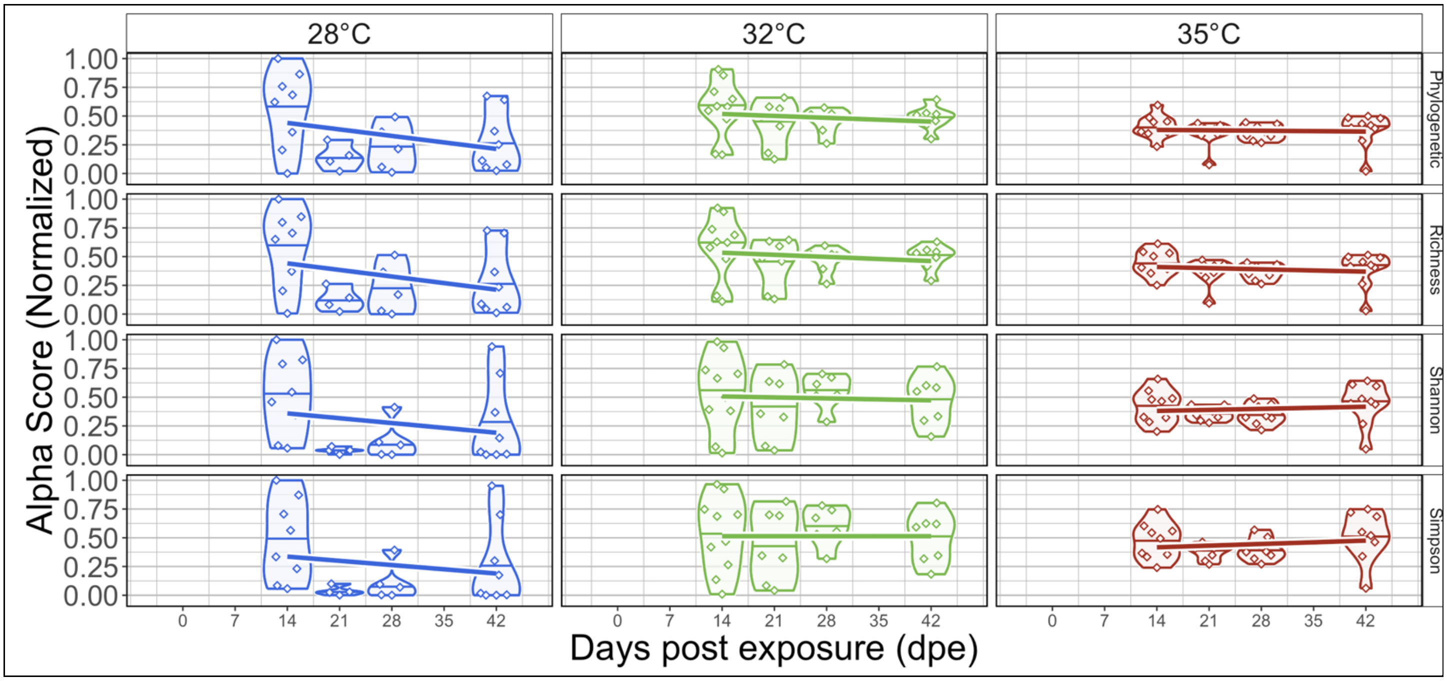
28°C
32°C
35°C
Phylogenetic
Richness
Shannon
Simpson
Alpha Score (Normalized)
1.00
0.75
0.50
0.25
0.00
0
7
14
21
28
35
42
Days post exposure (dpe)

S4D)

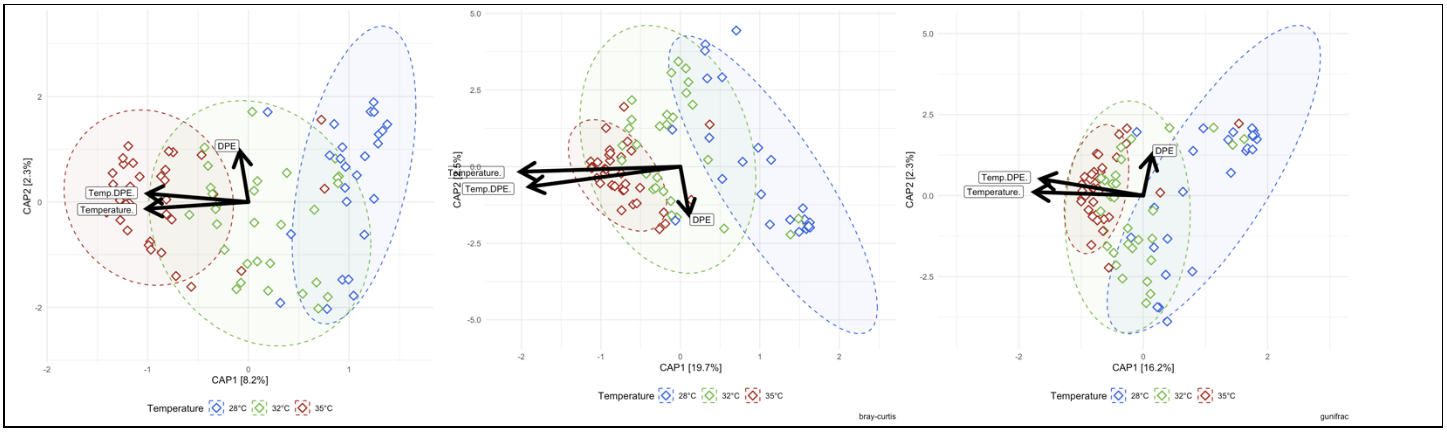
CAP2 [2.3%]
CAP1 [8.2%]
DPE
Temp.DPE.
Temperature.
Temperature 28°C 32°C 35°C
CAP1 [19.7%]
bray-curtis
CAP1 [16.2%]
CAP2 [2.3%]
gunifrac

### 5) Gut microbiome response has a non-linear relationship with infection burden

## S5A)

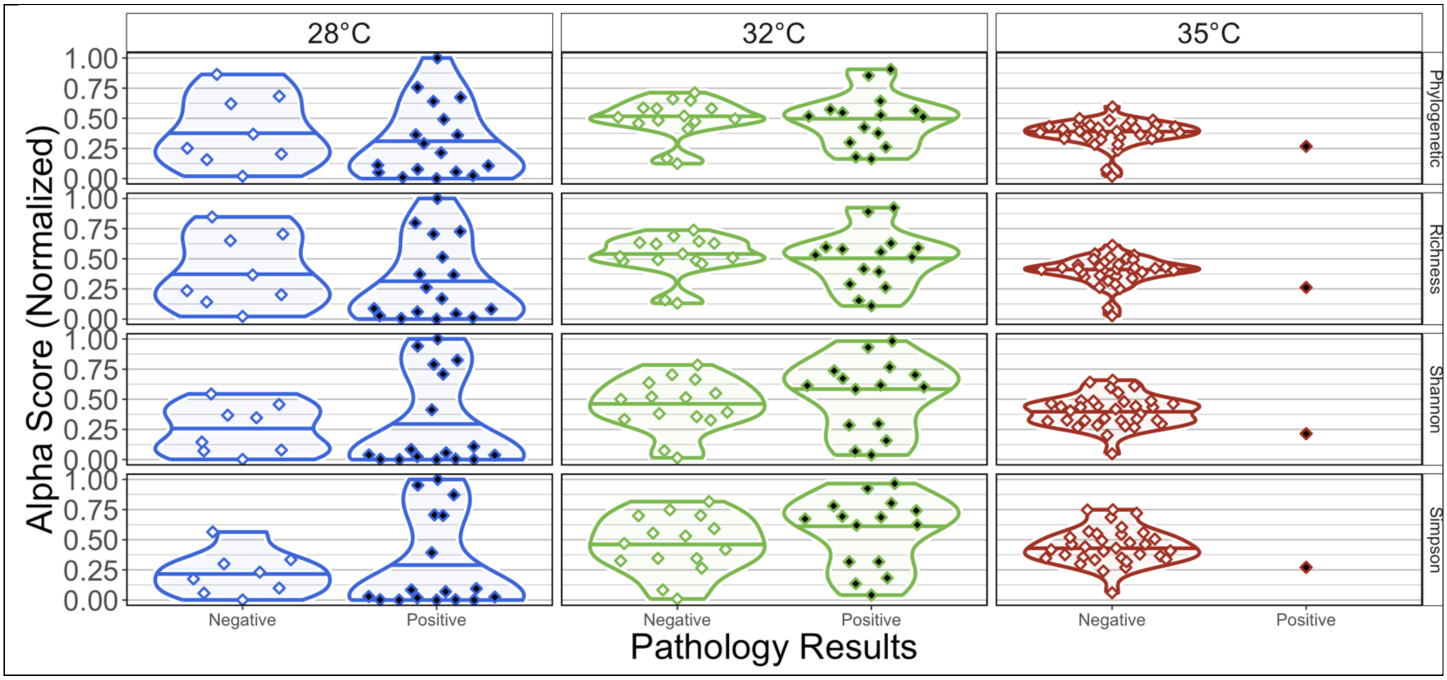

## S5B)

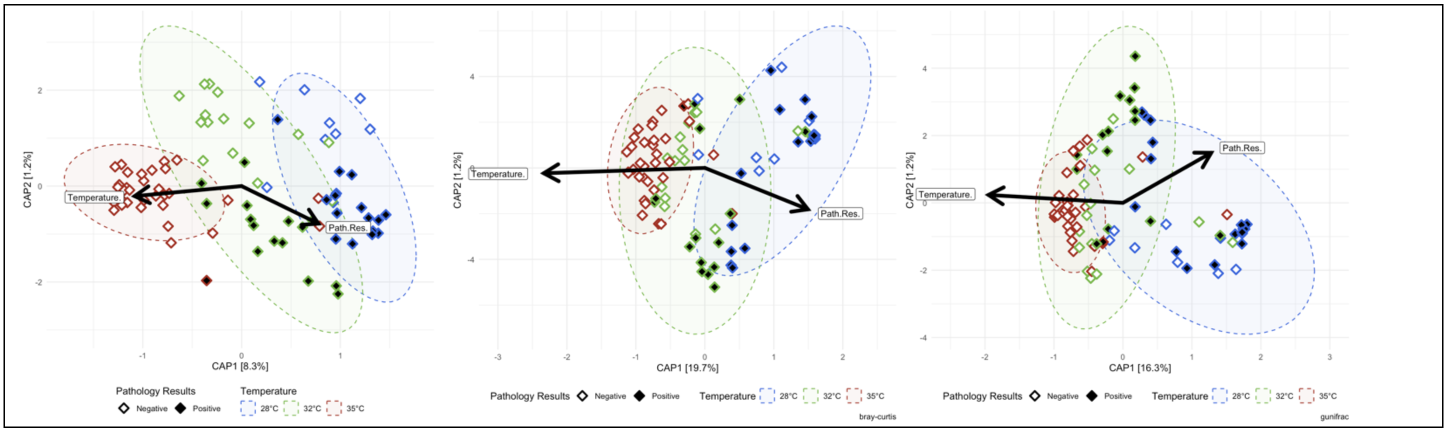

## S5C)

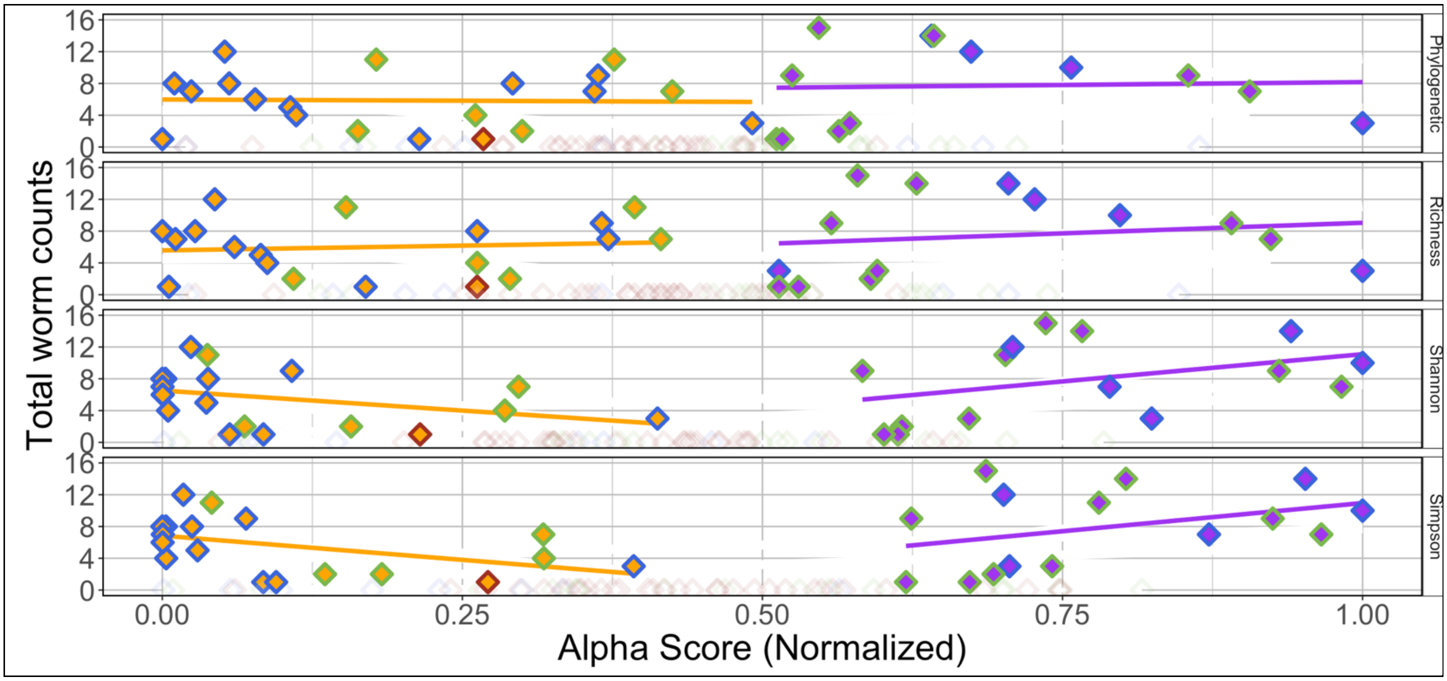
Total worm counts
Alpha Score (Normalized)
Phylogenetic
Richness
Shannon
Simpson
0.00
0.25
0.50
0.75
1.00

## S5D)

### S5D.1)

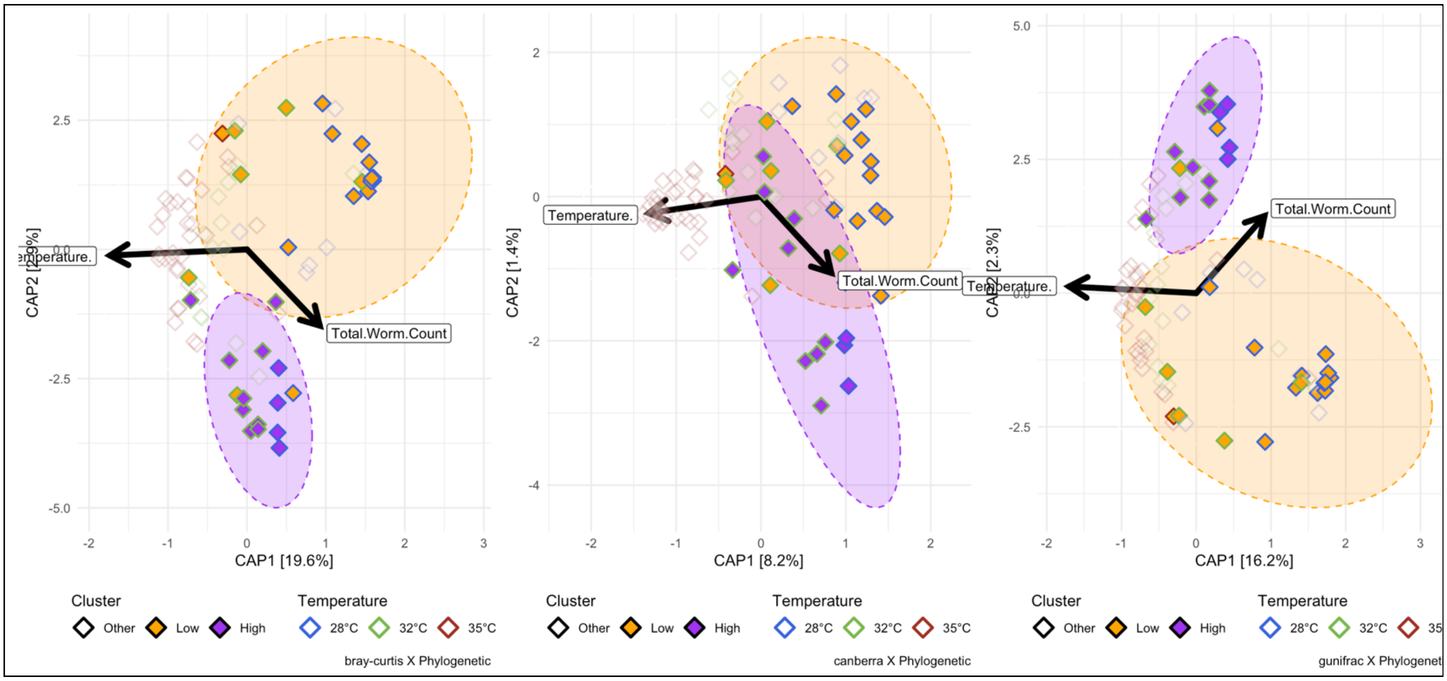
Total.Worm.Count
Temperature.
CAP1 [19.6%]
CAP1 [8.2%]
CAP2 [1.4%]
CAP1 [16.2%]
CAP2 [2.3%]
Cluster
Other
Low
High
Temperature
28°C
32°C
35°C
bray-curtis X Phylogenetic
canberra X Phylogenetic
gunifrac X Phylogenet

S5D.2)

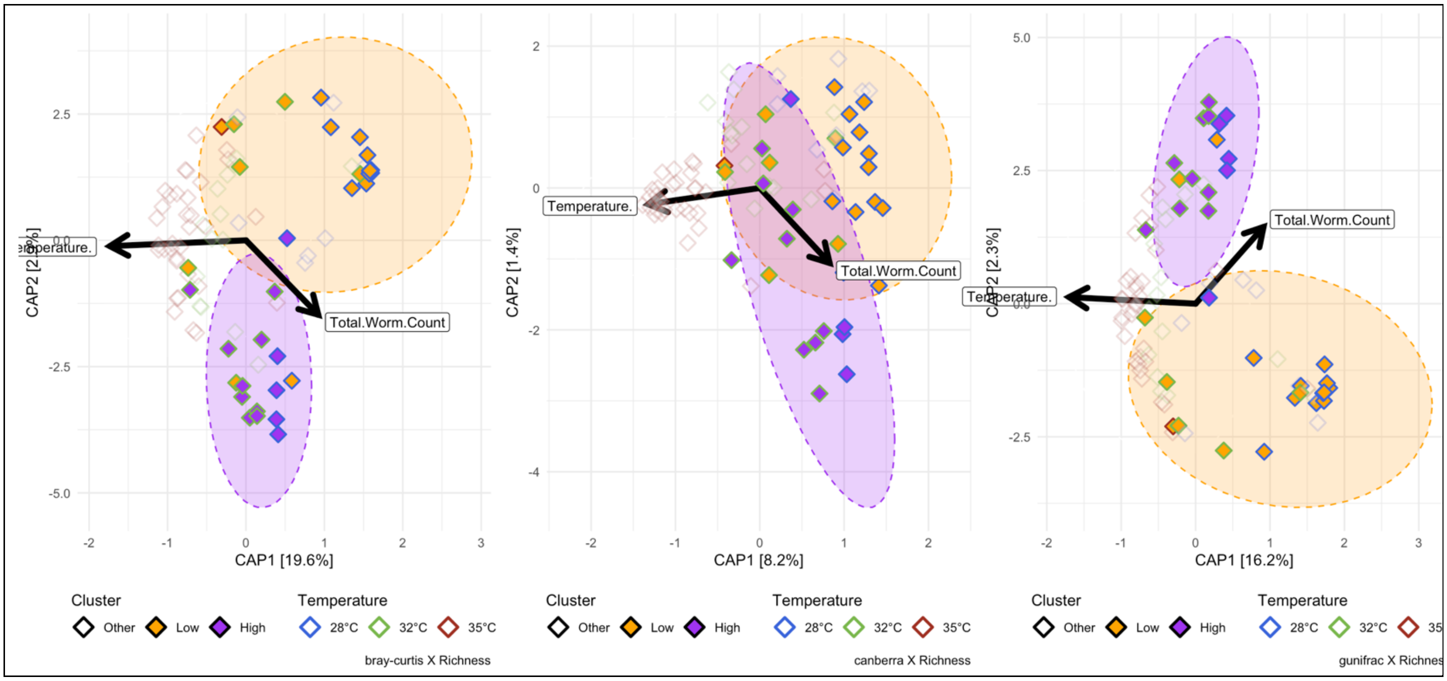

Temperature.
Total.Worm.Count
CAP1 [19.6%]
CAP1 [8.2%]
CAP1 [16.2%]
CAP2 [1.4%]
CAP2 [2.3%]
Cluster
Other
Low
High
Temperature
28°C
32°C
35°C
bray-curtis X Richness
canberra X Richness
gunifrac X Richnes

S5D.3)

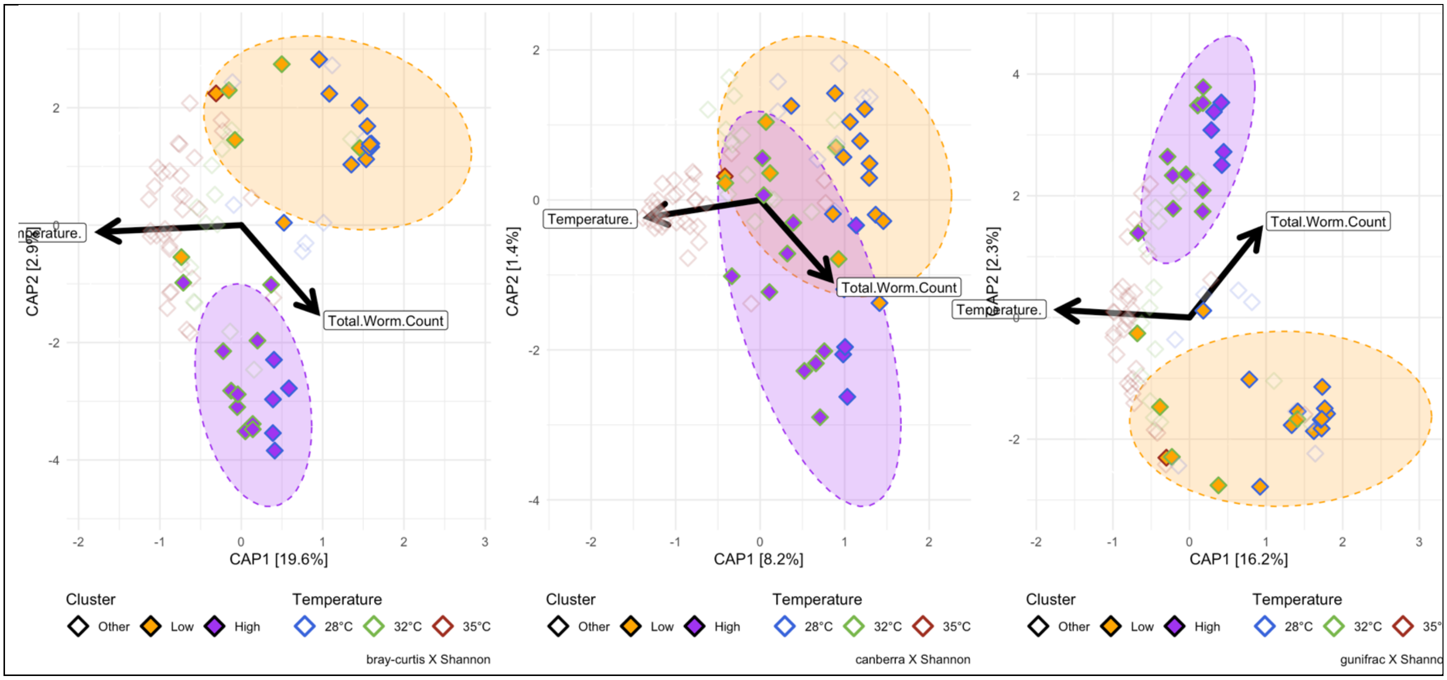

Temperature.
Total.Worm.Count
CAP1 [19.6%]
CAP1 [8.2%]
CAP1 [16.2%]
CAP2 [1.4%]
CAP2 [2.3%]
Cluster
Other
Low
High
Temperature
28°C
32°C
35°C
bray-curtis X Shannon
canberra X Shannon

S5D.4)

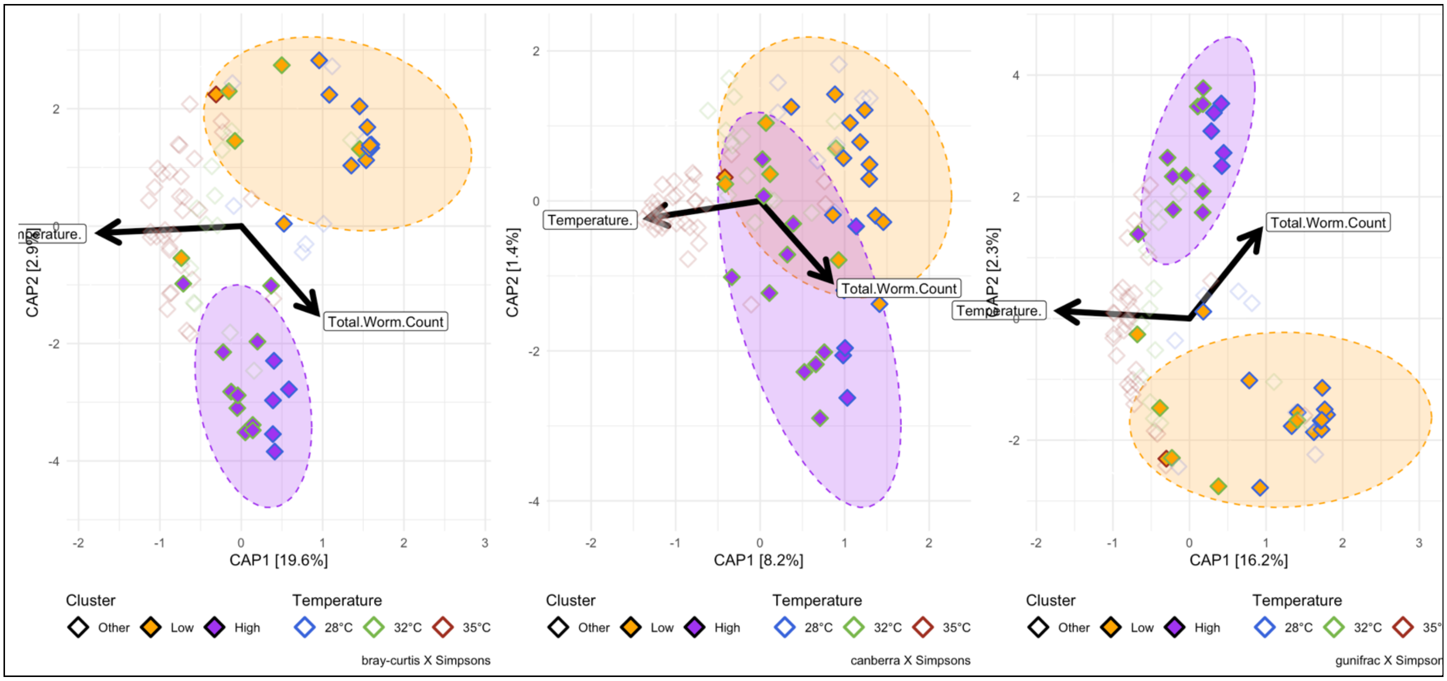

CAP2 [2.9%]
CAP1 [19.6%]
Temperature.
Total.Worm.Count
Cluster
Other
Low
High
Temperature
28°C
32°C
35°C
bray-curtis X Simpsons
CAP2 [1.4%]
CAP1 [8.2%]
Temperature.
Total.Worm.Count
Cluster
Other
Low
High
Temperature
28°C
32°C
35°C
canberra X Simpsons
CAP2 [2.3%]
CAP1 [16.2%]
Temperature.
Total.Worm.Count
Cluster
Other
Low
High
Temperature
28°C
32°C
gunifrac X Simpsor

#### 6) Parasite exposure exacerbates water temperature differences in gut microbiome structure

S6A)

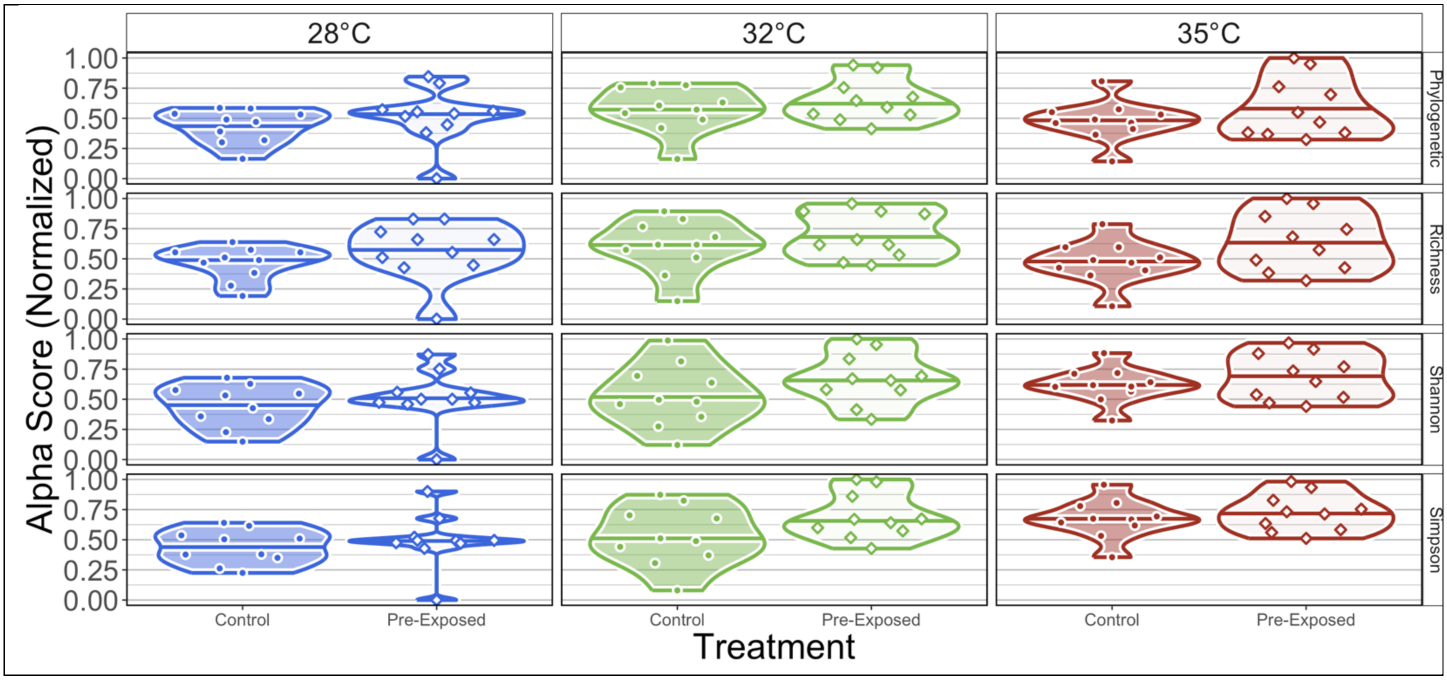

S6B)

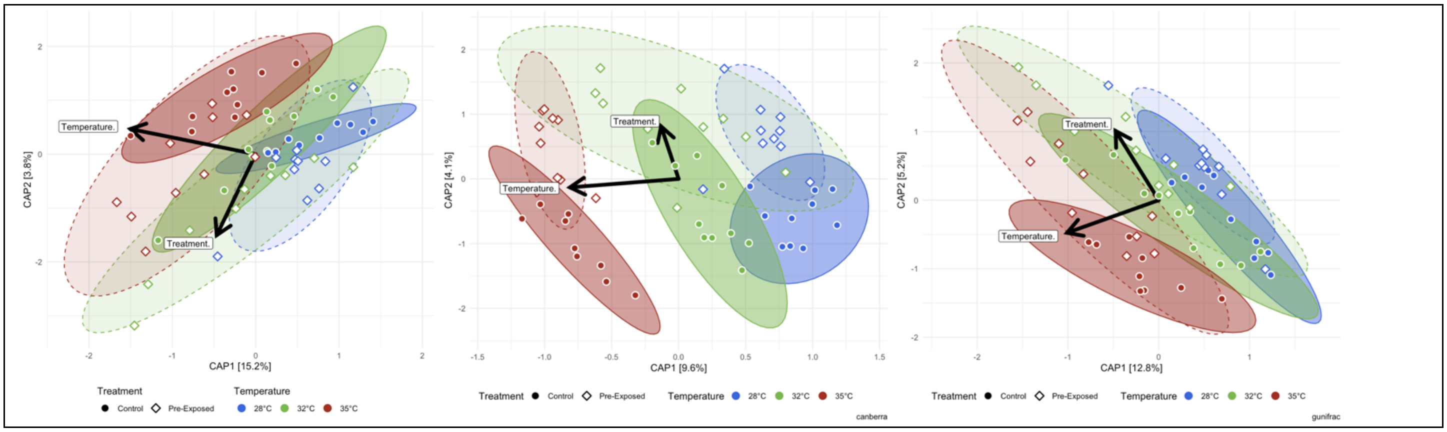

S6C)

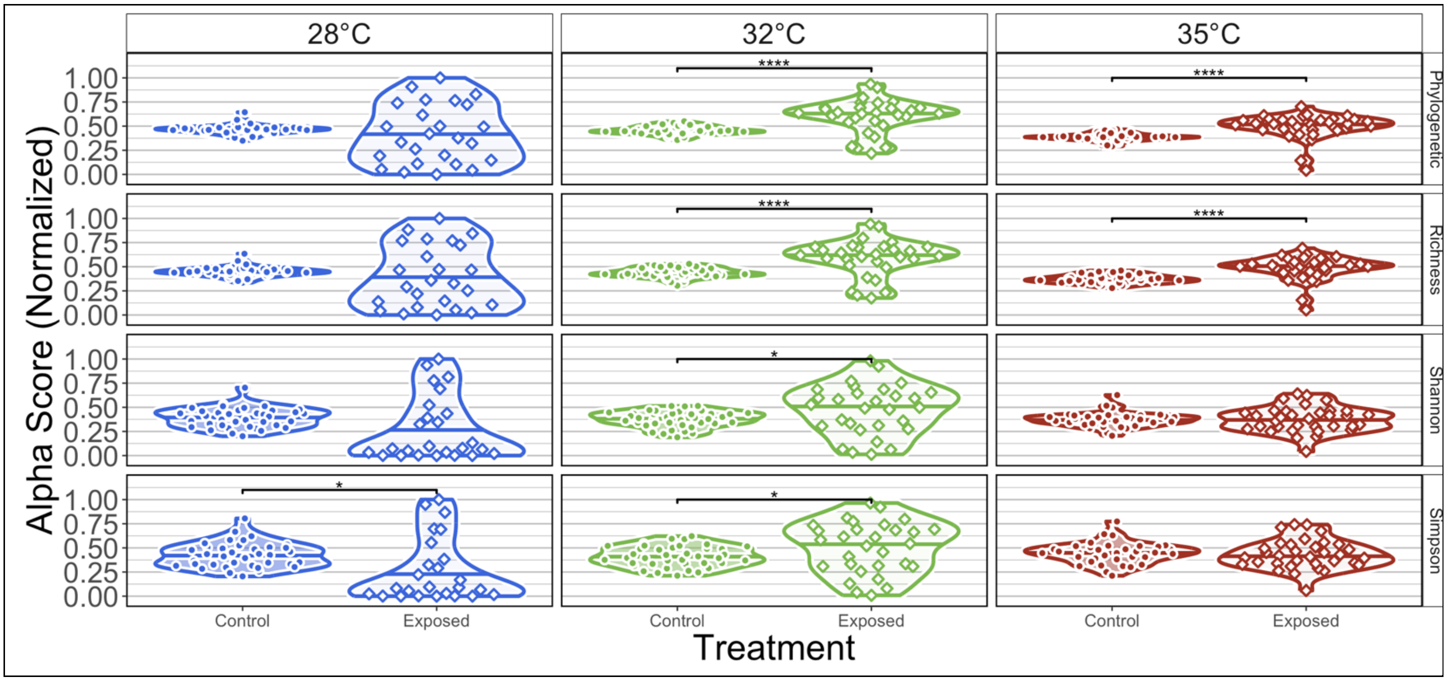
28°C
32°C
35°C
Phylogenetic
Richness
Shannon
Simpson
Alpha Score (Normalized)
1.00
0.75
0.50
0.25
0.00
****
*
Control
Exposed
Treatment

S6D)

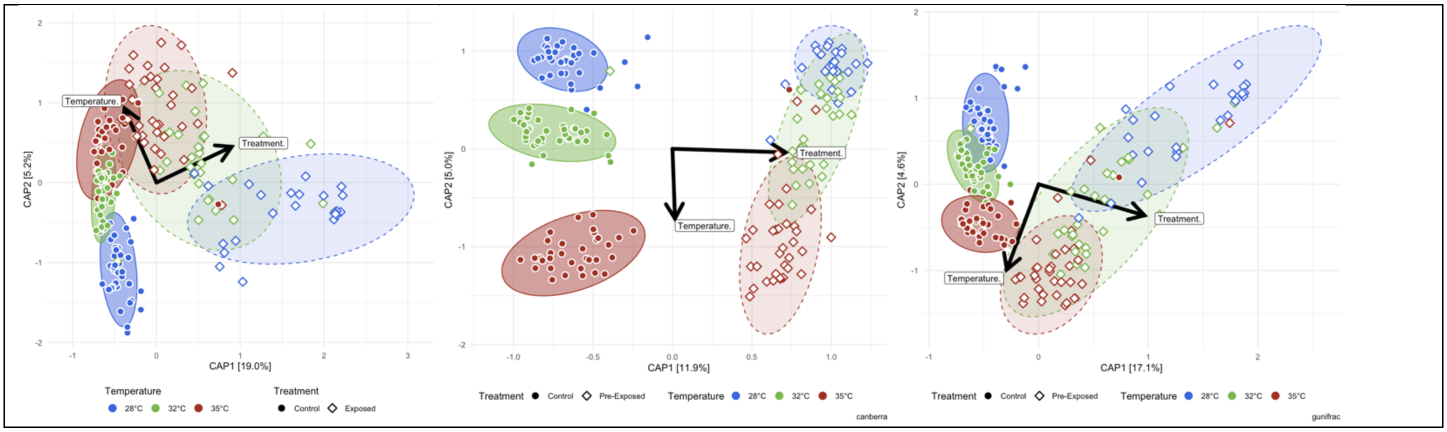
Temperature.
Treatment.
CAP1 [19.0%]
CAP2 [5.2%]
Temperature
28°C
32°C
35°C
Treatment
Control
Exposed
CAP1 [11.9%]
CAP2 [5.0%]
Pre-Exposed
canberra
CAP1 [17.1%]
CAP2 [4.6%]
gunifrac
